# Nrf2 Regulates β-cell Mass by Suppressing Cell Death and Promoting Proliferation

**DOI:** 10.1101/2021.03.05.434145

**Authors:** Sharon Baumel-Alterzon, Liora S. Katz, Gabriel Brill, Clairete Jean-Pierre, Yansui Li, Shyam Biswal, Adolfo Garcia-Ocaña, Donald K. Scott

## Abstract

Finding therapies that can protect and expand functional β-cell mass is a major goal of diabetes research. Here we generated β-cell-specific conditional knockout and gain-of-function mouse models and used human islet transplant experiments to examine how manipulating Nrf2 levels affects β-cell survival, proliferation and mass. Depletion of Nrf2 in β-cells resulted in decreased glucose-stimulated β-cell proliferation *ex vivo* and decreased adaptive β-cell proliferation and β-cell mass expansion after a high fat diet *in vivo*. Nrf2 protects β-cells from apoptosis after a high fat diet. Nrf2 loss-of-function decreases Pdx1 abundance and insulin content. Activating Nrf2 in a β-cell-specific manner increases β-cell proliferation and β-cell mass. Human islets transplanted under the kidney capsule of immunocompromised mice and treated systemically with CDDO-Me, an Nrf2 activator, display increased β-cell proliferation. Thus, Nrf2 regulates β-cell mass and is an exciting therapeutic target for expanding β-cell mass in diabetes.

## INTRODUCTION

One major characteristic of both major forms of diabetic patients is the lack of sufficient functional β-cell mass (Aguayo-Mazzucato and Bonner-Weir, 2018). Therefore, in order to therapeutically expand β-cell mass, several approaches have been explored including induced self-replication of β-cells and protection of existing β-cells (Aguayo-Mazzucato and Bonner-Weir, 2018). Oxidative stress is a major mediator of β-cell glucotoxicity and plays an essential role in the development of type 2 diabetes (T2D) (Gerber and Rutter, 2017; Newsholme et al., 2019; Yaribeygi et al., 2020). Indeed, chronic exposure of β-cells to high levels of reactive oxygen species (ROS) results in reduction of functional β-cell mass. If severe enough, ROS stimulates β-cell apoptosis through the mitochondrial “intrinsic” pathway by activation of the pro-apoptotic members of the Bcl-2 family and by recruitment of cytochrome C-mediated caspase 3 (Rojas et al., 2018). Increased ROS may also lead to defective glucose-stimulated insulin secretion (GSIS) in β-cells by inhibiting glycolytic enzymes, such as glyceraldehyde 3-phosphate dehydrogenase (Gapdh), pyruvate kinase M2 (Pkm2), and phosphofructokinase-1 (Pfk1), and by impairing the activity of potassium channels (Mullarky and Cantley, 2015; Yaribeygi et al., 2020). Additionally, glucotoxic ROS decrease the expression of the insulin gene by inhibiting the activity and levels of the duodenal homeobox factor 1 (Pdx1) and v-Maf musculoaponeurotic fibrosarcoma oncogene family, protein A (MafA) transcription factors (Robertson, 2006). Moreover, increased ROS activates the transcription factor FoxO1, which acts to inhibit β-cell proliferation under these conditions (Wang and Wang, 2017). Therefore, balancing ROS levels is critical for maintaining functional β-cell mass in diabetic patients.

Nuclear factor erythroid 2-related factor (Nrf2) is a well-studied transcription factor that confers cell protection against xenobiotic and oxidative stresses (Li et al., 2019; Panieri and Saso, 2019). During resting conditions, Nrf2 is bound to its repressor, the Kelch-like ECH-associated protein 1 (Keap1), which serves as an adaptor for the Cullin3-E3 ubiquitin ligase complex, thus targeting Nrf2 to proteosomal degradation. Upon oxidative stress, critical cysteines in Keap1 are oxidized, leading to disruption of Keap1-Keap1 homodimerization and Keap1-Cullin3 interaction, all of which disrupts Keap1’s ability to target Nrf2 for degradation. As a result, Nrf2 is phosphorylated at Ser40 and translocates to the nucleus where it binds to antioxidant response element (ARE) sequences in the regulatory regions of antioxidant target genes (Bloom and Jaiswal, 2003; Dinkova-Kostova et al., 2017b; Panieri and Saso, 2019; Suzuki and Yamamoto, 2017). Activation of Nrf2 has protective effects on β-cell survival and function, against a variety hazardous conditions, including damage induced by oxidative stress, streptozotocin, arsenic, glucotoxicity, glucolipotoxicity, cholesterol and cytokines (Baumel-Alterzon et al., 2020). In addition, Nrf2 preserves insulin content and insulin secretion, and regulates β-cell mitochondrial biogenesis and function (Baumel-Alterzon et al., 2020). These findings suggest that loss of Nrf2 function might play an important role in the development of diabetes. In agreement with this notion, GWAS analysis uncovered several mutations in the Nrf2 pathway that are associated with T2D (Jimenez-Osorio et al., 2016; Matana et al., 2020; Wang et al., 2015b). Recently, we found that Nrf2 stimulates β-cell proliferation in both rat insulinoma INS-1 cells and in primary pancreatic human islets (Kumar et al., 2018). We therefore hypothesized that activating Nrf2 preserves β-cell mass under diabetogenic-related stress conditions by increasing the survival of existing β-cells and promotes expansion of β-cell mass by stimulating β-proliferation.

Here, we show that Nrf2 is rapidly activated by glucose *in vitro*, and that Nrf2 is necessary for glucose-regulated β-cell proliferation. In addition, depletion of Nrf2 in an inducible β-cell-specific knockout mouse model results in impaired HFD-induced adaptive β-cell expansion due to increased β-cell death and decreased β-cell proliferation and insulin content. Conversely, genetic and pharmacological Nrf2 gain-of-function increases mouse β-cell proliferation and mass. Importantly, pharmacological Nrf2 gain-of-function stimulates human β-cell proliferation both *ex vivo* and *in vivo*. We conclude that the antioxidant Nrf2 pathway is an exciting novel target for increasing functional β-cell mass in diabetic patients.

## RESULTS

### Nrf2 is activated by glucose metabolism and ROS production in pancreatic β-cells

ROS are predominately generated by the mitochondria as a byproduct of increased glucose metabolism (Baumel-Alterzon et al., 2020). Cells exposed to high concentrations of glucose form ROS, which stimulate nuclear translocation of Nrf2 and activation of its transcriptional activity (Crean et al., 2012; Jiang et al., 2010). Once Nrf2 is released from Keap1 binding, it is phosphorylated at Ser40 (localized at the Keap1-binding site) and translocates to the nucleus (Bloom and Jaiswal, 2003; Bryan et al., 2013; Dinkova-Kostova et al., 2017b). Therefore, the use of Nrf2-ser40 (Nrf2-p) antibody (Fig. 1A) can be used to quantify the proportion of nuclear Nrf2 as a surrogate of Nrf2 activation. Surprisingly, to our knowledge, the effect of increased glucose metabolism on Nrf2 activation has never been studied in pancreatic β-cells. We found that exposure of the rat insulinoma INS-1 832/13 cell line with high (20 mM) glucose concentration for 5 min resulted in increased nuclear staining of Nrf2-p, similar to hydrogen peroxide (H_2_O_2_) (Supp. Fig. 1A). However, addition of the antioxidant, N-acetylcysteine (NAC), blocked Nrf2 nuclear recruitment, suggesting that glucose increases Nrf2 nuclear levels by ROS formation. The effect of high glucose on Nrf2 activity was validated by the increased expression (6-fold) of NAD(P)H quinone dehydrogenase 1 (Nqo1), a known Nrf2 target in β-cells (Kumar et al., 2018), in INS-1 832/13 cells that were incubated in high glucose medium for 6 h (Supp. Fig. 1B). To test if glucose metabolism has a similar effect in primary mouse β-cells, C57BL/6 mouse pancreatic islets were isolated, dispersed and incubated at 5.5 mM or 20 mM glucose. Similar to INS-1 832/13 cells, primary mouse β-cells displayed an increase in Nrf2 nuclear staining in just 5 min (Fig. 1B).

**Figure 1.**
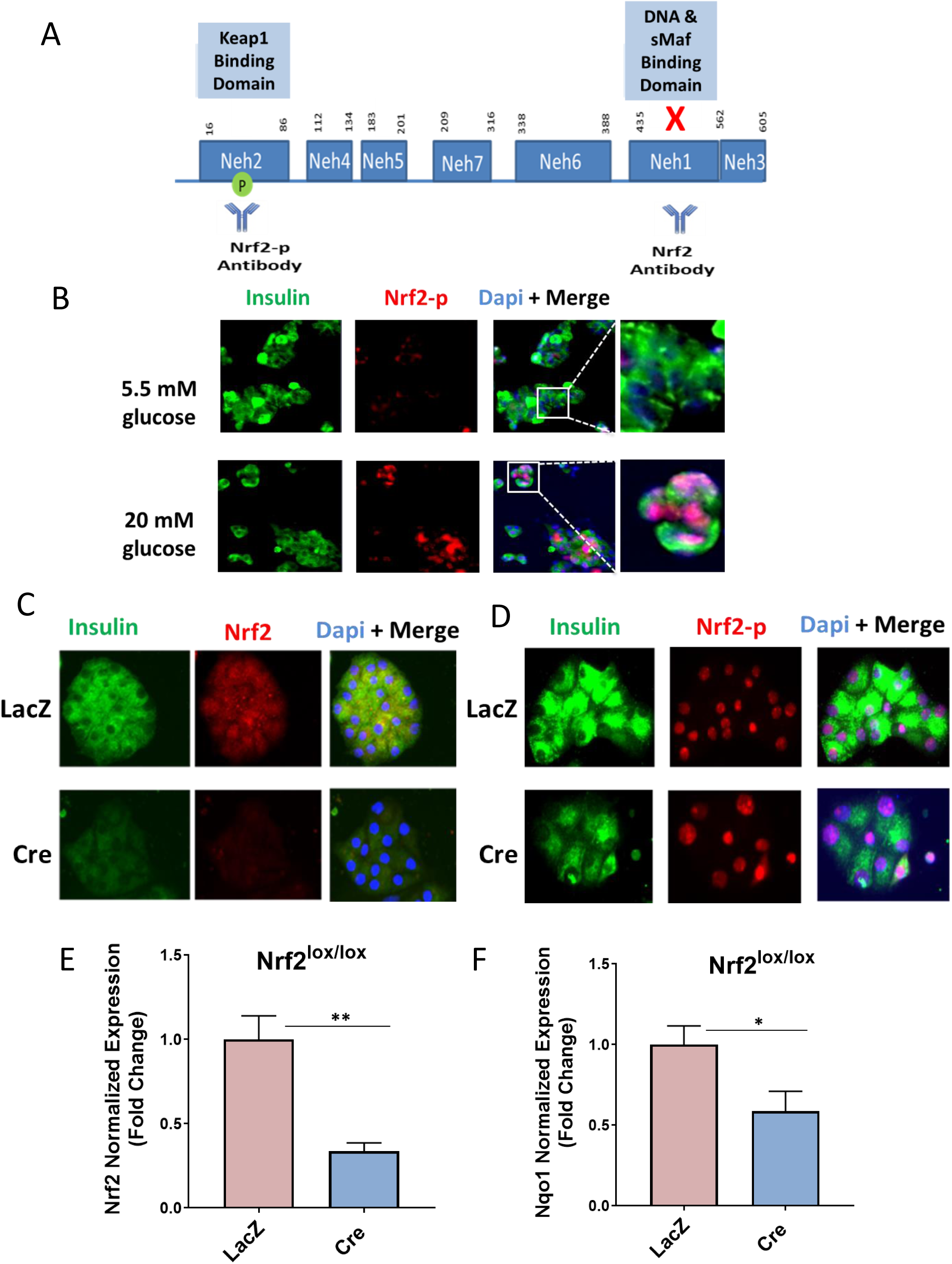
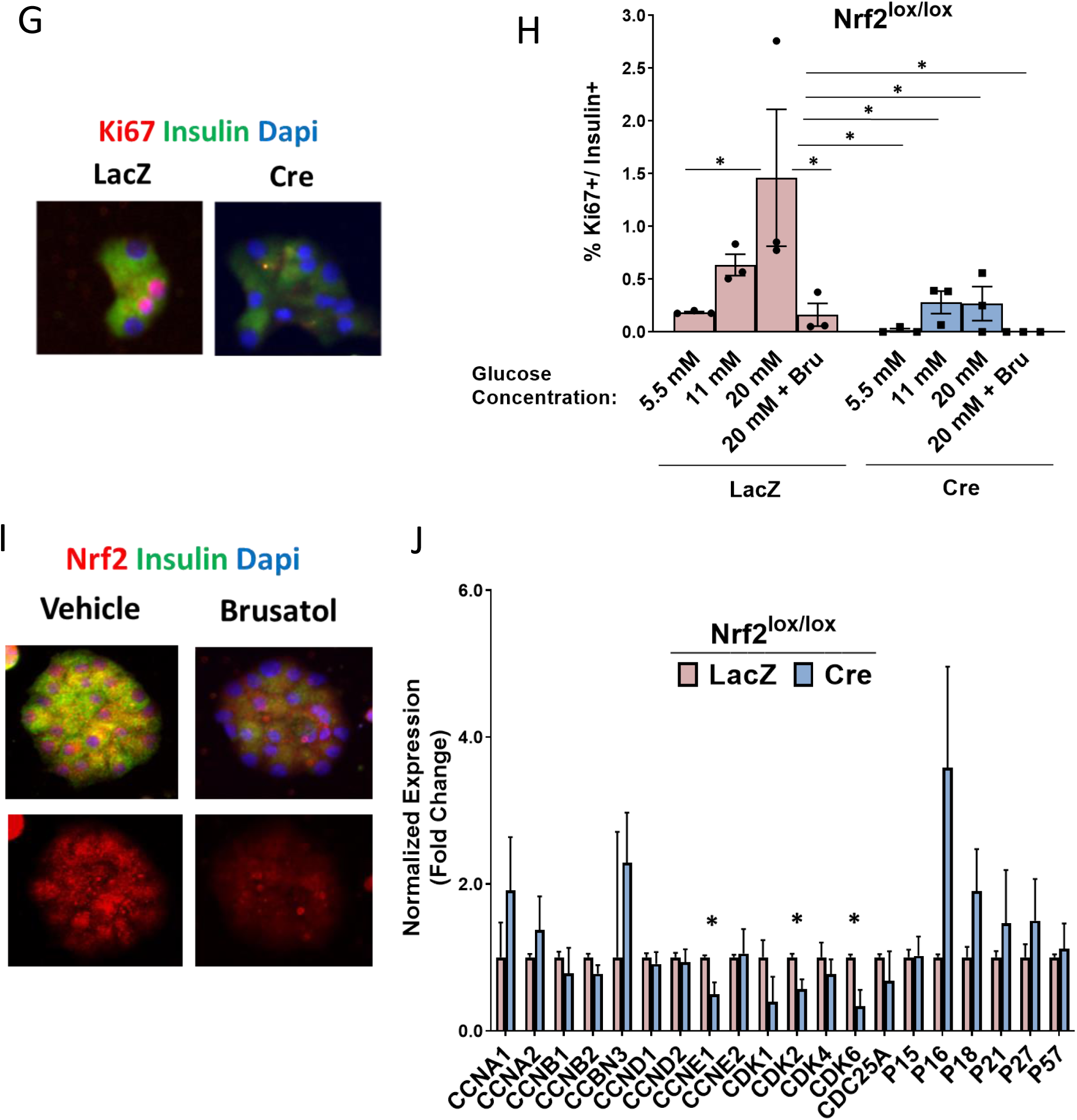
Nrf2 is necessary for glucose-stimulated β-cell proliferation *ex vivo.* (A) Illustration of Nrf2 protein structure emphasizing the deletion of exon 5, expressing the Neh1 domain, in Nrf2^lox/lox^ mice. The Nrf2 antibody (Nrf2 and Nrf2-p) epitopes are indicated. (B) Islets from C57Bl/6 mice were isolated, dispersed and incubated in 5.5 or 20 mM glucose for 5 min, followed by immunostaining with insulin and Nrf2-p antibodies. (C) Dispersed Nrf2^lox/lox^ mouse islets were transduced with LacZ or Cre adenovirus and cultured in the presence of 20 mM glucose for 72 h, followed by immunostaining using insulin and Nrf2 or (D) insulin and Nrf2-p antibodies. (E) Nrf2^lox/lox^ dispersed islets were incubated with 20 mM glucose for 72 h, followed by RNA isolation. Nrf2 mRNA expression using primers designed to identify wild-type transcripts (F) Nqo1 mRNA expression was measured to indicate Nrf2 activity. (G,H) Dispersed Nrf2^lox/lox^ mouse islets, transduced with LacZ or Cre adenovirus, and incubated with 50 nM Brusatol (Bru) or vehicle (100% ethanol) in the presence of 5.5, 11 or 20 mM glucose for 72 h, followed by immunostaining using insulin and Ki67 antibodies. Percent Ki67+ and Insulin+ cells was then calculated. (I) Mouse islets were incubated with 50 nM Brusatol or vehicle in the presence of 20 mM glucose for 72 h, followed by immunostaining with insulin and Nrf2 antibodies. (J) Nrf2^lox/lox^ islets were incubated in 20 mM glucose for 24 h, followed by RNA extraction. Expression of cell cycle regulators and β-cell identity genes was measured sing RT-PCR. Data shown are the mean +/-SEM (n=3-8, *p < 0.05; **p < 0.005).

### Nrf2 loss-of-function decreases glucose-stimulated β-cell proliferation *ex vivo*

We then tested if depletion of active Nrf2 affects glucose-stimulated β-cell proliferation. For this purpose, we used Nrf2^lox/lox^ mice (Reddy et al., 2011), in which Nrf2 exon 5 is flanked by LoxP sequences. Exon 5 encodes the Nrf2-ECH homology domain 1 (Neh1), necessary for DNA binding and for binding to sMaf, the Nrf2 obligatory heterodimer partner (Tonelli et al., 2018). Moreover, the Neh1 domain contains one (out of three) nuclear localization signal (NLS) and one (out of two) nuclear export signal (NES) (Giudice et al., 2010; Li et al., 2006; Theodore et al., 2008). Deletion of exon 5 in Nrf2 (confirmed by loss of fluorescence from an antibody that recognizes the Neh1 domain, Fig. 1A,C) did not prevent Nrf2 translocation into the nucleus (detected by Nrf2-p antibody, Fig 1A,D). However, the deletion of Nrf2 exon 5 led to an inactive product as we found a 70% decrease in Nrf2 expression as determined using an antibody recognizing the deleted region (Fig 1E), and a 50% reduction in the expression of its target gene, Nqo1 (Fig 1F). Glucose is a β-cell mitogen (Assmann et al., 2009; Gracia-ocana, 2010; Moulle et al., 2017; Stamateris et al., 2016). Previously, our lab has shown that Nrf2 overexpression stimulates β-cell proliferation in rat INS-1 832/13 cells and in primary human β-cells (Kumar et al., 2018). In order to test whether Nrf2 regulates the mitogenic effect of glucose in β-cells, isolated Nrf2^lox/lox^ islets were exposed to 5.5 mM, 11 mM or 20 mM glucose for 72 h. Incubation of control (LacZ) β-cells with high glucose concentration stimulated β-cell proliferation compared to incubation with 5.5 mM glucose (7.9-fold). However, this phenomenon was abolished in β-cells that expressed Cre, demonstrating that Nrf2 loss-of-function decreased glucose-stimulated β-cell proliferation (Fig. 1G,H). This result was confirmed using Brusatol, a natural quassinoid that blocks Nrf2 protein translation (Harder et al., 2017) (Fig. 1H,I and Supp. Fig. 1C,D).

In order to investigate the mechanism by which Nrf2 loss-of-function blocks glucose-stimulated β-cell proliferation, mouse Nrf2^lox/lox^ islets were isolated and cultured in low (5.5 mM) or high (20 mM) glucose. Following 24 h, cells were harvested, RNA was extracted and the expression of cell cycle regulators was measured. Depletion of Nrf2 decreased the expression of cyclin E1, a cyclin that regulates cell transition from G1 to S phase during cell division and is needed for DNA replication (Mazumder et al., 2004; Pils et al., 2014), and cyclin-dependent kinases 2 and 6 (CDK2 and CDK6), which are essential for cell cycle progression (Lim and Kaldis, 2013) (Fig. 1J).

### Nrf2 is necessary for HFD-mediated adaptive β-cell proliferation and β-cell mass expansion

Rats that are fed on a high fat diet (HFD) for 9 days display increased Nrf2 nuclear levels in β-cells and increased expression of Heme oxygenase 1, an Nrf2 target gene (Abebe et al., 2017), suggesting that the Nrf2 pathway is activated during overnutrition. To test whether the Nrf2 pathway is activated in mouse β-cells under similar conditions, mice were fed with a HFD or a regular diet (RD) for one week. As expected, mice that were fed with a HFD displayed increased Nrf2 levels in β-cells (3.3-fold) compared to RD-fed mice (Fig. 2A,B). Additionally, HFD-fed mice displayed increased expression of the Nrf2 target gene, Nqo1 (Fig. 2C), suggesting that the Nrf2 pathway is activated in mouse β-cells under hypercaloric conditions. Interestingly, we saw no change in Keap1 levels when comparing RD- and HFD-fed mice (Supp. Fig. 2A), suggesting that the activation of Nrf2 is not due to changes in Keap1 expression levels.

**Figure 2.**
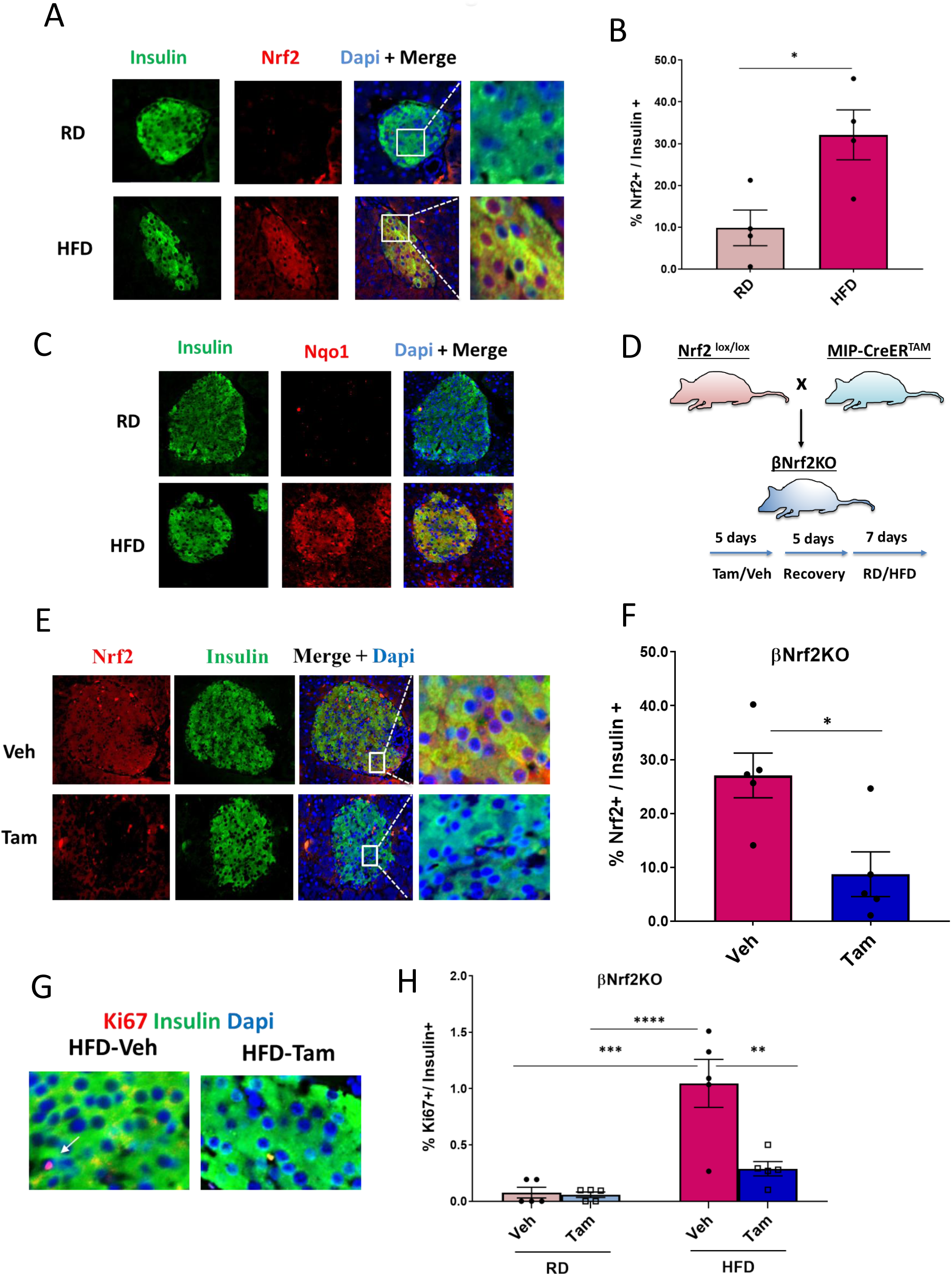
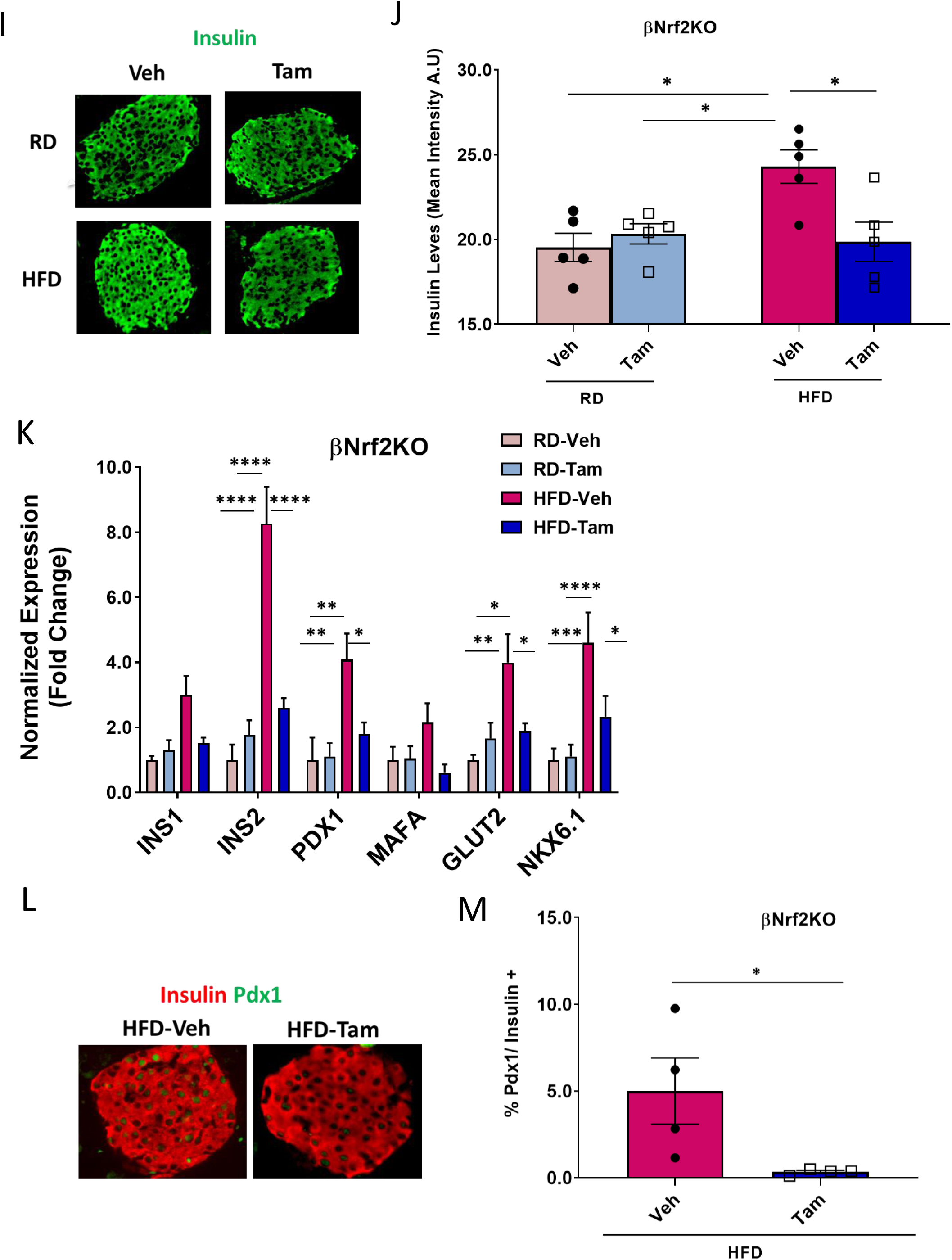
Nrf2 is necessary for HFD-adaptive β-cell proliferation *in vivo*. (A,B) C57Bl/6 mice were fed on HFD or RD for one week, after which their pancreata was removed, embedded and immunolabeled with insulin and Nrf2 antibodies. Percent Nrf2 nuclear positive/insulin positive cells were calculated. (C) Pancreata from C57Bl/6 mice fed on a RD or HFD for one week were immunolabeled with insulin and Nqo1 antibodies. (D) MIP-CreER^TAM^/Nrf2^lox/lox^ mice were crossed to generate βNrf2KO mice, which were injected daily for 5 days with Tam or Veh (corn oil), followed by 5 days of recovery and 1 week feeding with high fat diet (HFD) or regular diet (RD). (E,F) Pancreata from βNrf2KO mice fed on a HFD for one week immunolabeled with insulin and Nrf2 antibodies. Percent Nrf2 nuclear positive/insulin positive cells were calculated. (G,H) Pancreata from βNrf2KO mice fed a RD or HFD for one week immunolabeled with insulin and Ki67 antibodies. Percent proliferation in insulin positive cells was then calculated. (I,J) Pancreata from βNrf2KO mice fed on a HFD or RD for one week immunolabeled with insulin. Mean intensity of insulin positive cells was calculated. (K) Islets were isolated from βNrf2KO mice fed on a HFD or RD for one week, RNA was extracted and expression of β-cell identity genes was measured. (L,M) Pancreata from βNrf2KO mice fed on a HFD for one week immunolabeled with insulin and Pdx1 antibodies. Percent Pdx1 nuclear positive/insulin positive cells were calculated. Data shown are the mean +/-SEM (n=4-6, *p < 0.05; **p < 0.005, ***p < 0.0005, ****p < 0.0001).

Mice fed one-week on a HFD have increased β-cell proliferation as an adaptive response to increased insulin demand (Mosser et al., 2015; Stamateris et al., 2013). To test the role of Nrf2 in HFD-adaptive β-cell proliferation, we established a tamoxifen-inducible β-cell-specific Nrf2 loss-of-function mouse model by crossing MIP-CreER^TAM^ mice (Wicksteed et al., 2010) with Nrf2^lox/lox^ mice (Reddy et al., 2011), named here βNrf2KO mice. βNrf2KO mice were injected with corn oil vehicle or tamoxifen, followed by a recovery period, after which they were fed for one-week with RD or HFD (Fig. 2D). Immunostaining of the pancreata from HFD-fed βNrf2KO mice using Nrf2 and insulin antibodies (Fig. 2E,F), show that tamoxifen-injected mice had a 60% reduction in Nrf2 levels in β-cells compared to vehicle controls when both are fed on HFD, thus confirming the deletion of Nrf2 exon 5 in β-cells. As expected, feeding of corn oil-injected mice on a HFD for one week resulted in stimulation of β-cell proliferation compared to RD-fed mice (8-fold) (Fig. 2G,H). A similar effect was also observed in MIP-CreER^TAM^ control mice (Supp. Fig. 2B). However, this adaptive increase of β-cell proliferation was abolished in tamoxifen-injected βNrf2KO mice (Fig. 2G,H), demonstrating that Nrf2 is necessary for HFD-stimulated β-cell proliferation.

Our results showed that both the genetic and the pharmacological Nrf2 loss-of-function in β-cells under metabolic stress resulted in reduced insulin staining intensity compared to control (Fig. 1C,D,G,I and Supp. Fig. 3A,B), raising the possibility that Nrf2 depletion affects β-cell insulin content. In order to test this hypothesis, mouse Nrf2^lox/lox^ islets were isolated and exposed to high (20 mM) glucose concentration. Following 72 h, cells were harvested, and insulin content was measured. Cre-expressing Nrf2^lox/lox^ islets had a 40% reduction in insulin content (Supp. Fig. 3C). Concordantly, tamoxifen-injected βNrf2KO mice, which were fed on HFD for one week presented with a 20% decrease in insulin staining, compared to vehicle control mice (Fig. 2I,J). The same results were found using a different source of insulin antibody (data not shown). Based on these findings, we next analyzed the expression of β-cell identity genes in these βNrf2KO islets. There was a significant reduction in the expression of Ins2, Pdx1, Glut2 and Nkx6.1 in tamoxifen-injected βNrf2KO mice compared to vehicle controls when fed on HFD (Fig. 2K). Reduced levels of Pdx1 were also observed in pancreata after depletion of Nrf2 (Fig. 2L,M). In Nrf2^lox/lox^ islets, a reduction in Pdx1 and MafA expression was observed in Cre expressing islets compared to vehicle treated control (Supp. Fig. 3D). Thus, deletion of Nrf2 in β-cells under metabolic stress leads to decreased insulin expression and content.

**Figure 3.**
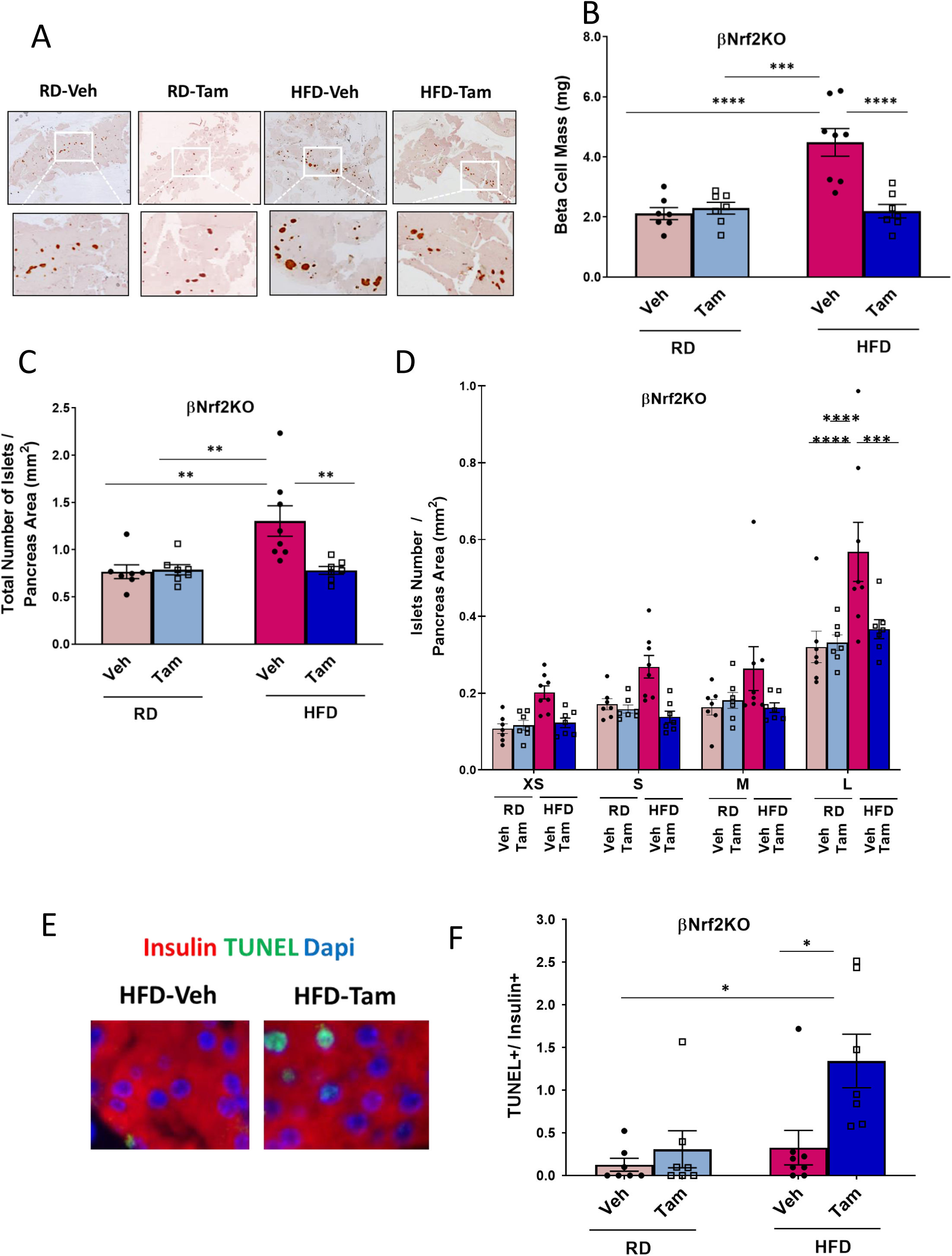
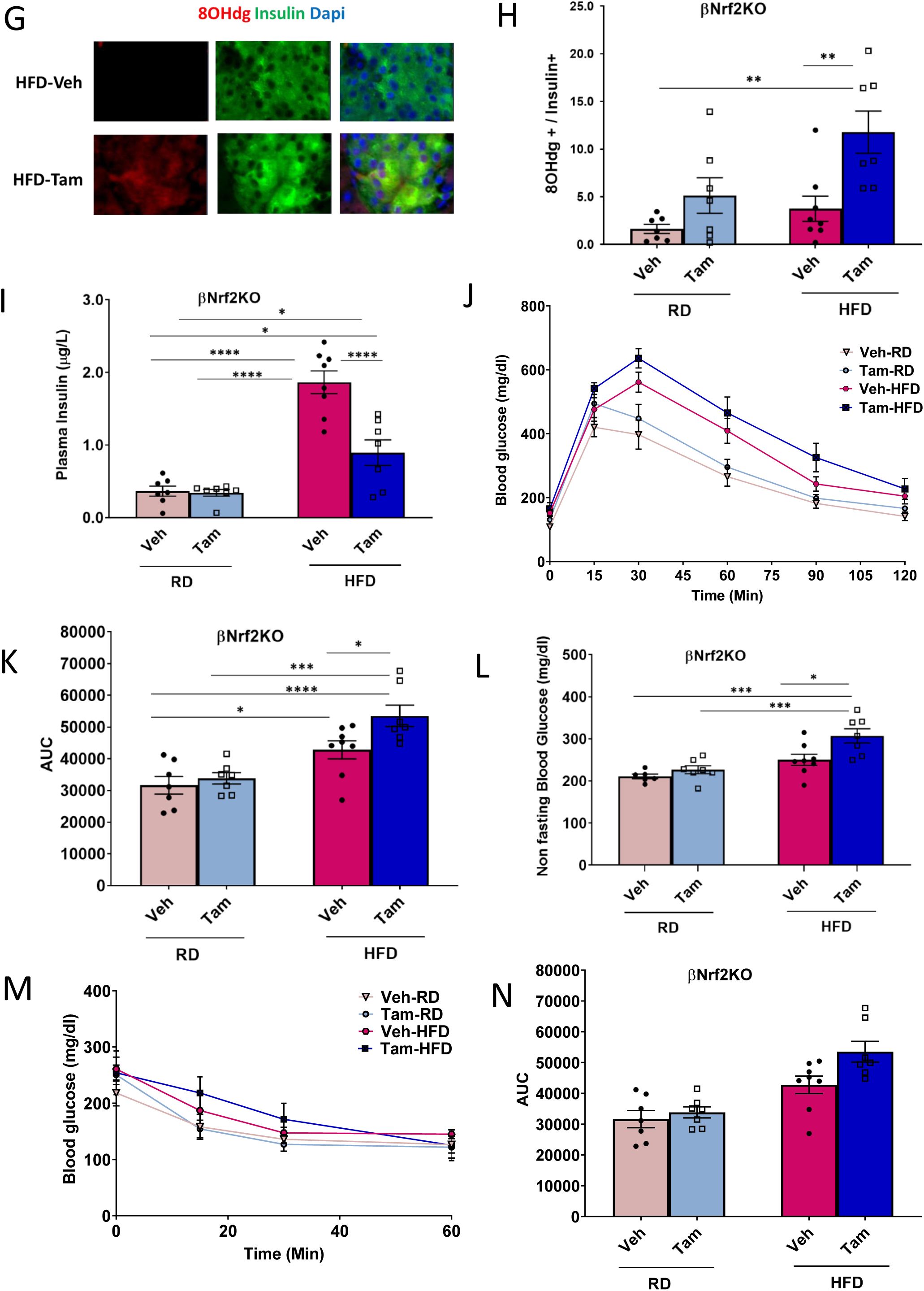
Nrf2 is necessary for HFD-mediated β-cell mass expansion. Pancreata from βNrf2KO mice fed a RD or HFD for 29 days were immunolabeled for insulin. β-cell mass (A,B) and islet morphometry was calculated. (C) Total islet number per pancreas area. (D) Islet size was divided to four groups XS (<1000 µm^2^), S (1001-2200 µm^2^), M (2201-4400 µm^2^), L (>4400 µm^2^) and islet numbers per pancreas area calculated. (E,F) The percentage of apoptotic insulin positive cells was calculated using a TUNEL assay in pancreata from HFD-fed βNrf2KO mice. (G,H) Pancreata from βNrf2KO mice were immunolabeled for insulin and 8-OHdG oxidative stress marker. 8-OHdG nuclear positive/insulin positive cells were calculated. (I) Plasma insulin was measured from βNrf2KO mice following 29 days of HFD or RD feeding. (J,K) Intraperitoneal glucose tolerance test (ipGTT) performed in βNrf2KO mice after an overnight fast. AUC was calculated. (L) Non-fasting blood glucose. (M,N) Insulin sensitivity test (ITT) performed in βNrf2KO mice in the fed state after 29 days on a RD or HFD. AUC was calculated. Data shown are the mean +/-SEM (n=7-8, *p < 0.05; **p < 0.005, ***p < 0.0005, ****p < 0.0001).

Adaptive proliferation in response to a HFD leads to significantly increased β-cell mass after 2-4 weeks, and since β-cell proliferation is a major contributor to the adaptive expansion of β-cell mass (Meier et al., 2008), we tested if depletion of Nrf2 affects adaptive β-cell mass expansion after 1 month on a HFD. Vehicle or tamoxifen-injected βNrf2KO mice were placed on a HFD or RD for 28 days, pancreata were harvested and immunostained with an insulin antibody for β-cell mass analysis (Fig. 3A,B). As expected, HFD-fed control mice displayed an adaptive increase in β-cell mass (2.3-fold) compared to mice that were fed on RD. However, this adaptive increase was not observed in HFD-fed tamoxifen-injected mice, suggesting that depletion of Nrf2 impaired adaptive expansion of β-cell mass induced by HFD. HFD-fed control mice displayed an increase in total islet number (1.7-fold) compared to RD-fed controls; however, this increase was abolished in HFD-fed tamoxifen-injected βNrf2KO mice (Fig. 3C). Furthermore, Nrf2-depleted mice displayed a 35% decrease in the largest islets, with no significant changes in the number of smaller islets, in mice fed on HFD compared to controls (Fig. 3D). Thus, Nrf2 is required for the expansion of larger islets after a HFD.

Nrf2 promotes β-cell survival under various stresses (Baumel-Alterzon et al., 2020). Therefore, we investigated whether Nrf2 loss-of-function attenuates HFD-adaptive β-cell expansion by decreasing β-cell survival. Tamoxifen-injected βNrf2KO mice on a HFD had 18.8-fold more TUNEL positive β-cells compared to control mice on a HFD (Fig. 3E,F). In addition, immunostaining using an antibody against the oxidative damage marker, 8-OHdG, showed a 6.2-fold increase in oxidative stress in β-cells depleted of Nrf2 compared to control mice fed a HFD [Fig. 3G,H, (Valavanidis et al., 2009)]. Thus, the presence of Nrf2 protects β-cells from the increased oxidative stress and cell death elicited by a HFD.

As expected, due to increased insulin demand, vehicle control βNrf2KO mice that were fed on HFD had increased plasma insulin levels (5.1-fold) compared to mice fed on RD (Fig. 3I). However, the increase in insulin levels was abolished in HFD-fed tamoxifen-injected βNrf2KO mice. Accordingly, a significant reduction in glucose tolerance (Fig. 3J,K) and an increase in non-fasting blood glucose levels (Fig. 3L) were observed in tamoxifen-injected mice compared control mice fed a HFD. No significant changes in body weight or fasting blood glucose were observed in Nrf2-depleted mice compared to control mice on a HFD (Supp. Fig. 4A,B). Additionally, no significant changes in insulin sensitivity were observed (Fig. 3M,N), indicating that the effects on glucose tolerance were not due to defective insulin signaling. Thus, Nrf2 is necessary for maintaining glucose homeostasis after a HFD.

**Figure 4.**
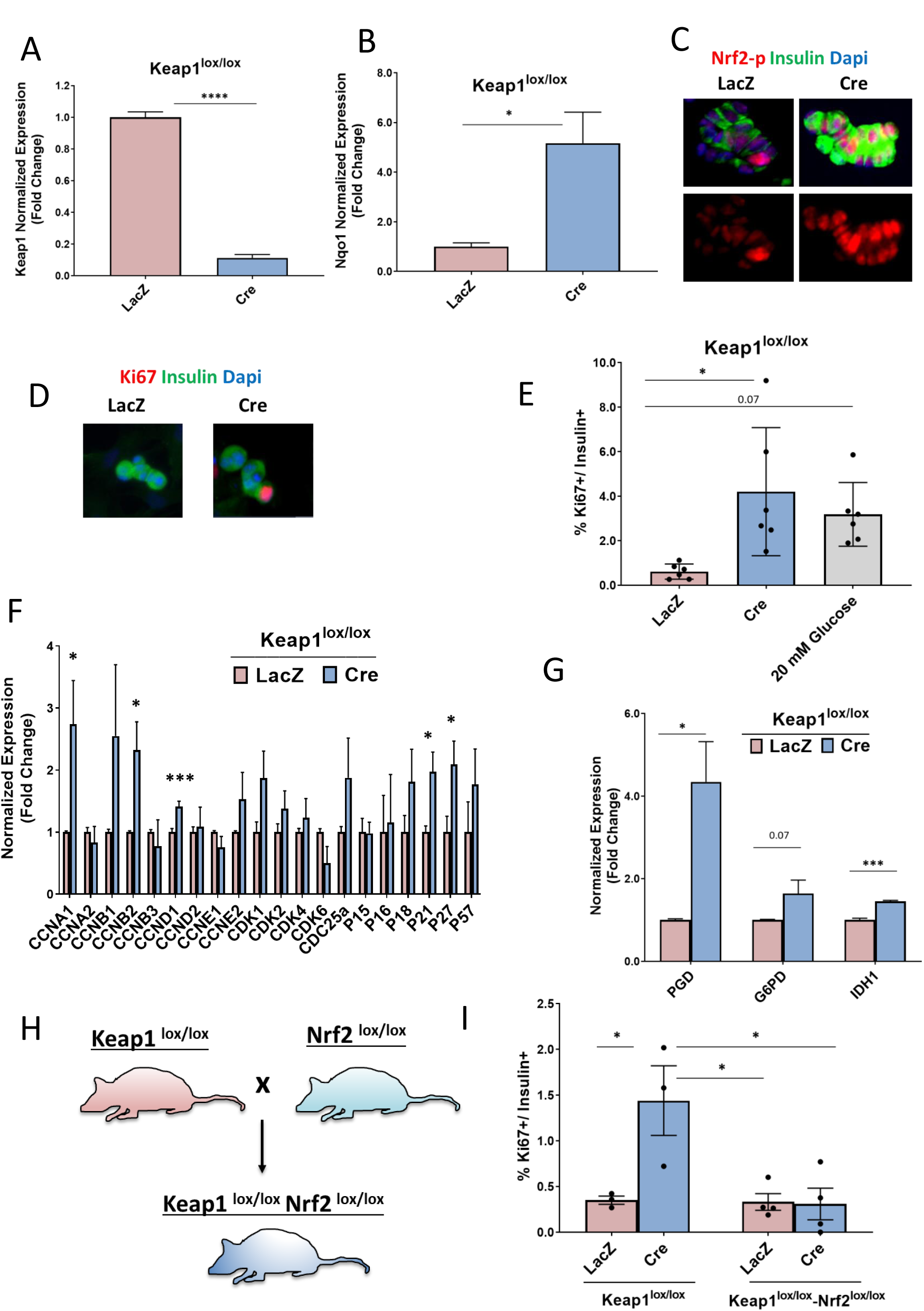
Genetic Nrf2 gain-of-function increases β-cell proliferation *ex vivo*. Dispersed Keap1^lox/lox^ islets were transduced with Cre or LacZ expressing adenoviruses. Following 72 h, RNA was isolated and Keap1 (A) or Nqo1 mRNA expression (B) was measured. (C) Keap1^lox/lox^ islets were immunolabeled with insulin and Nrf2-p, or insulin and Ki67 antibodies (D). The percentage of proliferating insulin positive cells was calculated (E). (F,G) Keap1^lox/lox^ islets were isolated and transduced with Cre or LacZ adenoviruses. After 24 h, RNA was extracted and the expression of cell cycle regulators (F) and Nrf2 target genes (G) was measured. (H,I) Keap1^lox/lox^ mice were crossed with Keap1^lox/lox^-Nrf2^lox/lox^ to make double knockout mice. Nrf2^lox/lox^ Keap1^lox/lox^-Nrf2^lox/lox^ or Keap1^lox/lox^ islets were isolated, transduced with Cre or LacZ adenoviruses, and stained for insulin and Ki67. The percentage of Ki67 positive and insulin positive cells was calculated. Data shown are the mean +/-SEM (n=3-8, *p < 0.05; ***p < 0.0005).

### Nrf2 gain-of-function increases β-cell proliferation *ex vivo*

Since depletion of Nrf2 leads to decreased glucose and HFD-stimulated β-cell proliferation we next tested if Nrf2 gain-of-function stimulates β-cell proliferation. Under normal physiological conditions, Nrf2 levels are regulated by its inhibitor, Keap1, which targets Nrf2 to proteosomal degradation (Dinkova-Kostova et al., 2017b). Therefore, to increase Nrf2 levels, we used Keap1 ^lox/lox^ mice, in which LoxP sites flank Keap1 exons 2 and 3. In this model, Cre-mediated recombination results in a nonfunctional Keap1 protein that lacks the redox-sensitive intervening region (IVR) domain and the Nrf2-binding domain (Blake et al., 2010). Islets from Keap1^lox/lox^ were isolated, cultured in 5.5 mM glucose, and transduced with adenoviruses expressing either LacZ as a control or Cre recombinase. Cre-expressing islets had a 90% decrease in Keap1 RNA expression (Fig. 4A) concomitant with 5.1-fold increase in RNA expression of the Nrf2 target gene, Nqo1 (Fig. 4B), and increased levels of nuclear pSer40 Nrf2 (Fig. 4C), consistent with Nrf2 activation. The same islets displayed a nearly 7-fold increase in β-cell proliferation, comparable to that induced by 20 mM glucose (Fig. 4D,E).

To investigate further, RNA was extracted from LacZ-or Cre-transduced Keap1^lox/lox^ islets and expression of cell cycle regulators was measured. Keap1 deletion significantly increased expression of cyclins A1, B2, and D1, in concordance with increased β-cell proliferation. Somewhat surprisingly, Keap1 deletion also significantly increased expression of the cell cycle inhibitors p27 and p21 (Fig. 4F), though similar effects have been reported in other cell types (Chung et al., 2015; Li et al., 2011; Malhotra et al., 2010). No significant changes in the expression of β-cell maturity genes were observed (Supp. Fig 5A). Nrf2 supports cell proliferation by controlling expression of genes that are involved in generation of nicotinamide adenine dinucleotide phosphate (NADPH) and in purine nucleotide synthesis, both needed for dividing cells (Fu et al., 2019; Hayes and Ashford, 2012; Mitsuishi et al., 2012). These genes include glucose-6-phosphate dehydrogenase (G6pd) and phosphogluconate dehydrogenase (Pgd) of the pentose phosphate pathway, and isocitrate dehydrogenase 1 (Idh1). We found that depletion of Keap1 increased expression of Pgd (4.3-fold increase), and Idh1 (1.4-fold increase) (Fig 4G), suggesting that Nrf2 supports β-cell proliferation in part by increasing NADPH levels.

**Figure 5.**
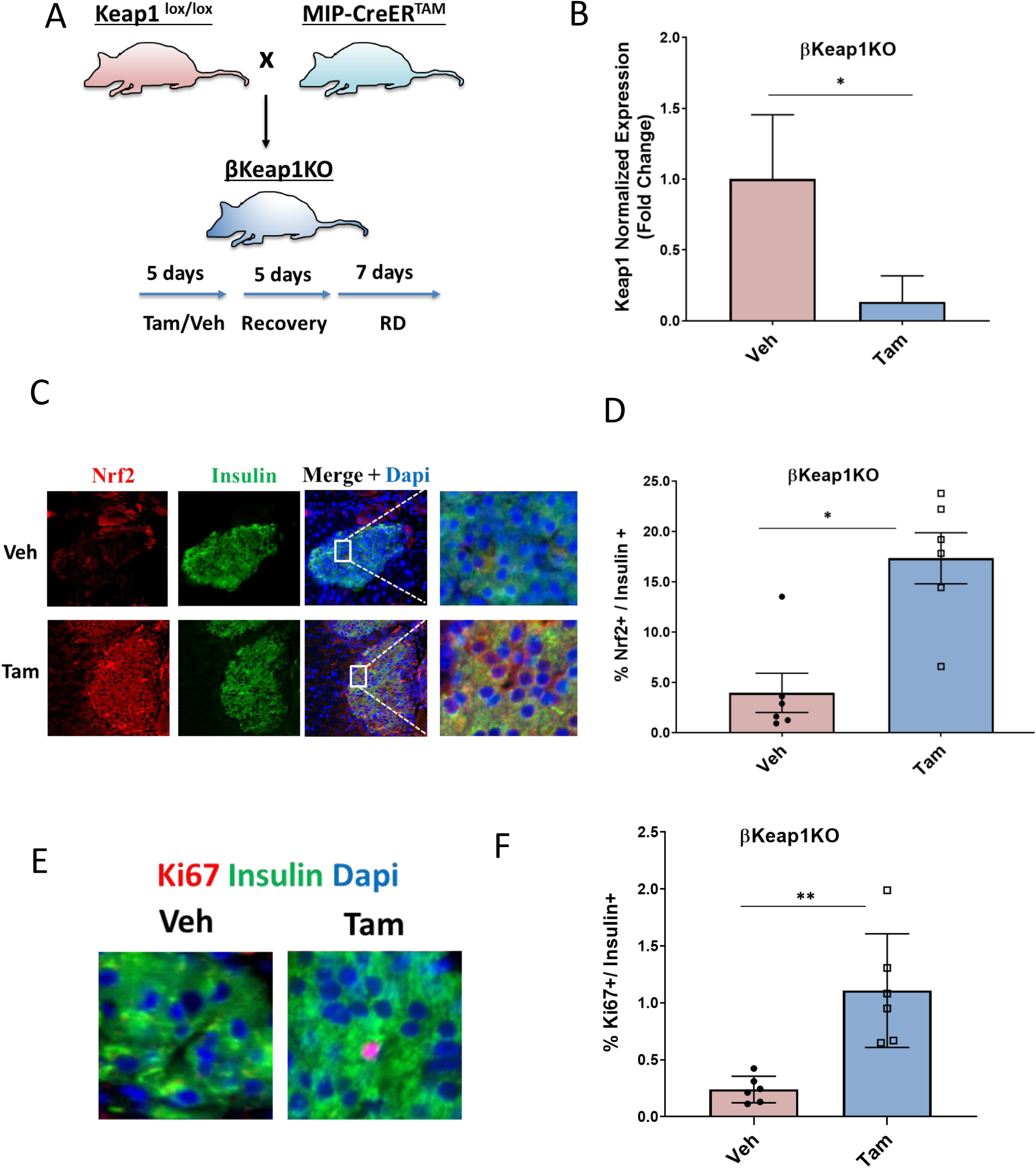
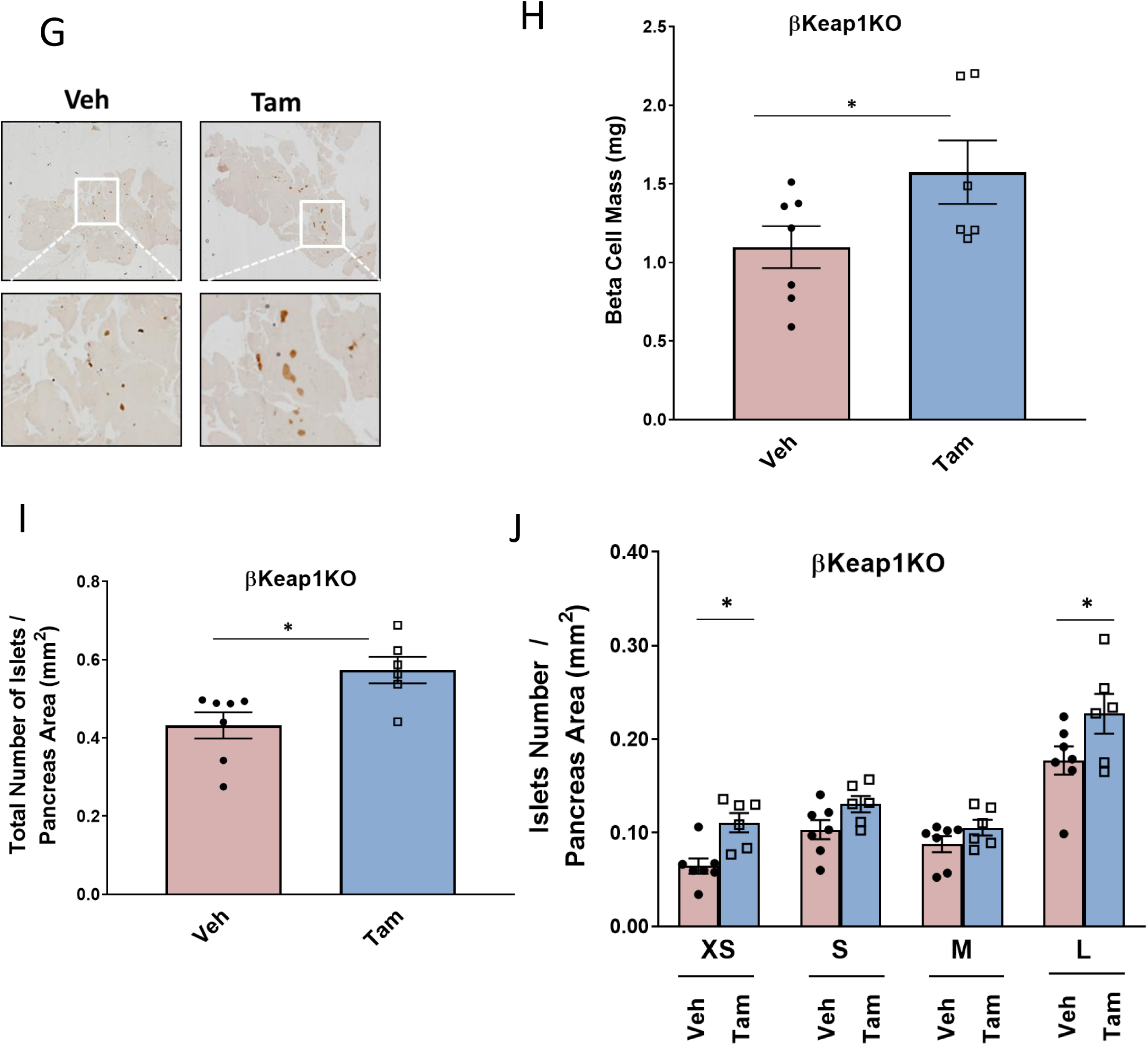
Genetic Nrf2 gain-of-function increased β-cell proliferation and mass *in vivo*. (A) Mip-CreER^TAM^ and Keap1^lox/lox^ mice were crossed to generate βKeap1KO mice, which were injected daily for 5 days with Tam or Veh (corn oil), followed by 5 days of recovery and 1 week on a regular diet (RD). (B) Islets were isolated from βKeap1KO mice, RNA was extracted and Keap1 expression was measured. (C,D) Pancreata from βKeap1KO mice were removed, embedded and immunolabeled with insulin and Nrf2 antibodies. The percentage Nrf2 nuclear events in insulin positive cells were calculated. (E,F) Pancreata from βKeap1KO were immunolabeled with insulin and Ki67 antibodies, and the percent Ki67 positive/insulin positive cells was calculated. (G-J) βKeap1KO pancreata from Tam- or Veh-treated mice were immunolabeled with insulin. β-cell mass and islet morphometry (total islets number per pancreatic area) were calculated. Data shown are the mean +/-SEM (n=3-7, *p < 0.05; **p < 0.005).

Although Keap1 is considered to be the canonical inhibitor of Nrf2, several other proteins bind Keap1 (Komatsu et al., 2010; Lo and Hannink, 2008; Tamberg et al., 2018; Wan et al., 2020). Therefore, in order to test whether the increased proliferation induced by Keap1 deletion is specific to Nrf2 activation, we generated double knockout mice (Keap1^lox/lox^-Nrf2^lox/lox^) in which both Keap1 and Nrf2 can be deleted by Cre recombinase (Fig. 4H). As expected, deletion of Keap1 resulted in increased β-cell proliferation (4.1-fold), but there was no increased proliferation when both Keap1 and Nrf2 were deleted (Fig. 4I). Thus, deletion of Keap1 increases β-cell proliferation due to activation of the Nrf2 pathway.

### Nrf2 gain-of-function *in vivo* stimulates β-cell proliferation and increases β-cell mass

We then generated conditional β-cell-specific Keap1 knockout mice by crossing MIP-CreER^TAM^ mice with Keap^lox/lox^ mice to obtain MIP-CreER^TAM^/Keap1^lox/lox^, or βKeap1KO mice (Blake et al., 2010; Wicksteed et al., 2010) Fig. 5A]. Deletion of Keap1 in tamoxifen-injected mice resulted in an 80% decrease of Keap1 mRNA and increased β-cell Nrf2 level 4.4-fold compared to control mice (Fig. 5B-D). Tamoxifen-injected βKeap1KO mice had a 4.6-fold increase in β-cell proliferation compared to control mice (Fig. 5E,F). Thus, increasing Nrf2 abundance stimulates β-cell proliferation both *ex vivo* and *in vivo*.

To test whether Nrf2 gain-of-function *in vivo* increases β-cell mass, pancreata from βKeap1KO mice were immunostained with an insulin antibody to perform β-cell histomorphometry. Deletion of Keap1 for I month resulted in increased β-cell mass (1.5-fold), increased total number of islets (1.3-fold) and increased the number of extra-small (1.7-fold) and large (1.3-fold) islets (Fig. 5G-J). These findings indicate that Nrf2 gain-of-function increases β-cell mass by increasing β-cell proliferation and increasing the number of islets. However, no significant changes in glucose homeostasis, body weight or plasma insulin were observed in βKeap1KO mice under basal conditions (Supp. Fig.6A-H).

**Figure 6.**
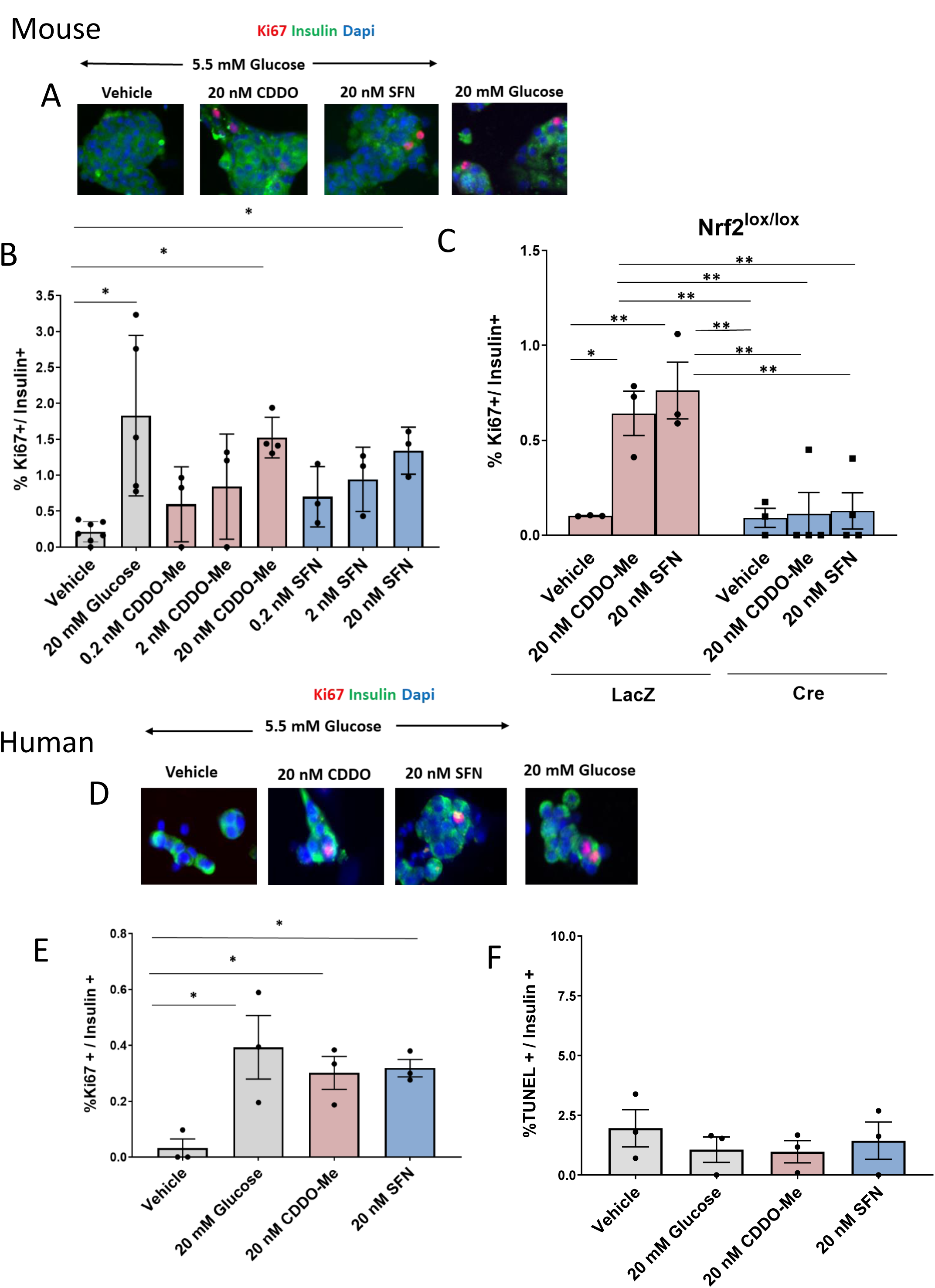
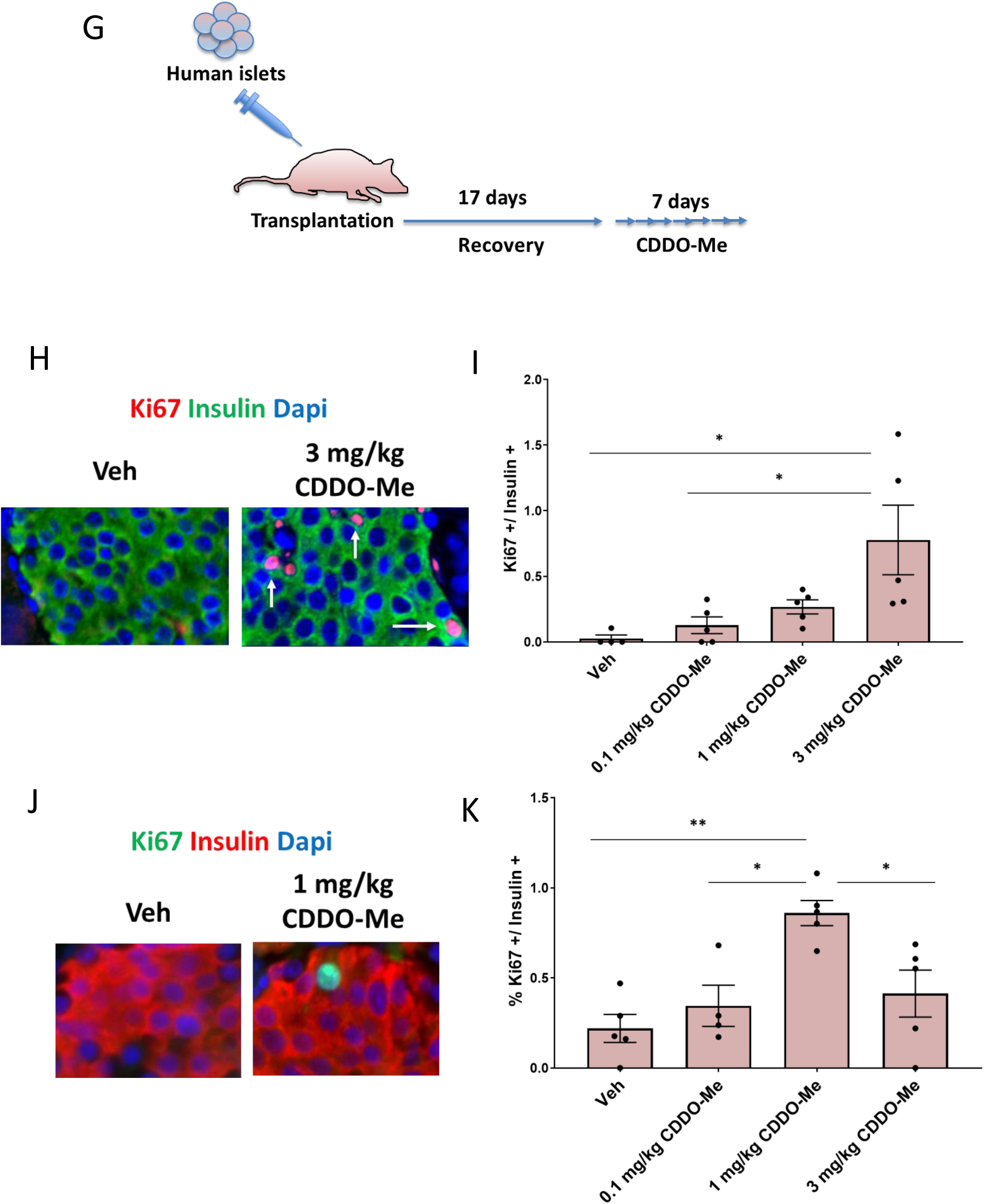
Pharmacological gain of Nrf2 function increases mouse and human β-cell proliferation. (A,B) C57BL/6 mouse islets were isolated, dispersed and incubated with increasing doses of CDDO-Me or SFN for 72 h, followed by immunostaining with insulin and Ki67 antibodies. The percentage of Ki67 and insulin doubly-positive cells were calculated. Islet cells treated with 20 mM glucose were used as a positive control. (C) Dispersed Nrf2^lox/lox^ mouse islets were incubated with 20 nM CDDO-Me or 20 nM SFN for 72 hours followed by immunostaining using insulin and Ki67 antibodies. Percent proliferation in insulin positives cells was calculated. (D,E) Human islets were incubated with 20 nM CDDO-Me or 20 nM SFN for 72 h followed by immunostaining with insulin and Ki67 antibodies, and the percentage of insulin and Ki67 positive cells was calculated. (F) A TUNEL assay was performed on human islets treated as shown. (G) Immunodeficient mice were transplanted with 500 human islets under the kidney capsule. Following a 17-day recovery, daily IP-injections were made with the indicated doses of CDDO-Me for 1 week. The pancreata (H,I) and kidney grafts (J,K) were immunostained with insulin and Ki67, and the percentage of insulin and Ki67 positive cells was calculated. Data shown are the mean +/-SEM (n=3-5, *p < 0.05; **p < 0.005).

### Pharmacological stimulation of Nrf2 increases β-cell proliferation

The synthetic triterpenoid bardoxolone methyl, a C-28 methyl ester of 2-cyano-3,12-dioxoolean-1,9-dien-28-oic acid (CDDO-Me), inhibits Keap1:Nrf2 interaction, and activates the Nrf2 antioxidant pathway (Cleasby et al., 2014; Sova and Saso, 2018). CDDO-Me is currently being tested in clinical trials for treating several conditions such as chronic kidney disease in diabetic patients, Alport syndrome and pulmonary hypertension (Baumel-Alterzon et al., 2020; Robledinos-Anton et al., 2019; Toto, 2018). Members of the Bardoxolone drug family activate Nrf2 and increase functional β-cell mass in mice by improving β-cell viabillty and function (Baumel-Alterzon et al., 2020). Interestingly, CDDO-Me time- and dose-dependently increased INS-1 832/13 cell number compared to vehicle control (Supp. Fig 7A). Additionally, 20 nM CDDO-Me increased Nrf2-p immunostaining in INS1 cells following 50 min of incubation, which validates the activation of Nrf2 under these conditions (Supp.Fig 7B).

**Figure 7.**
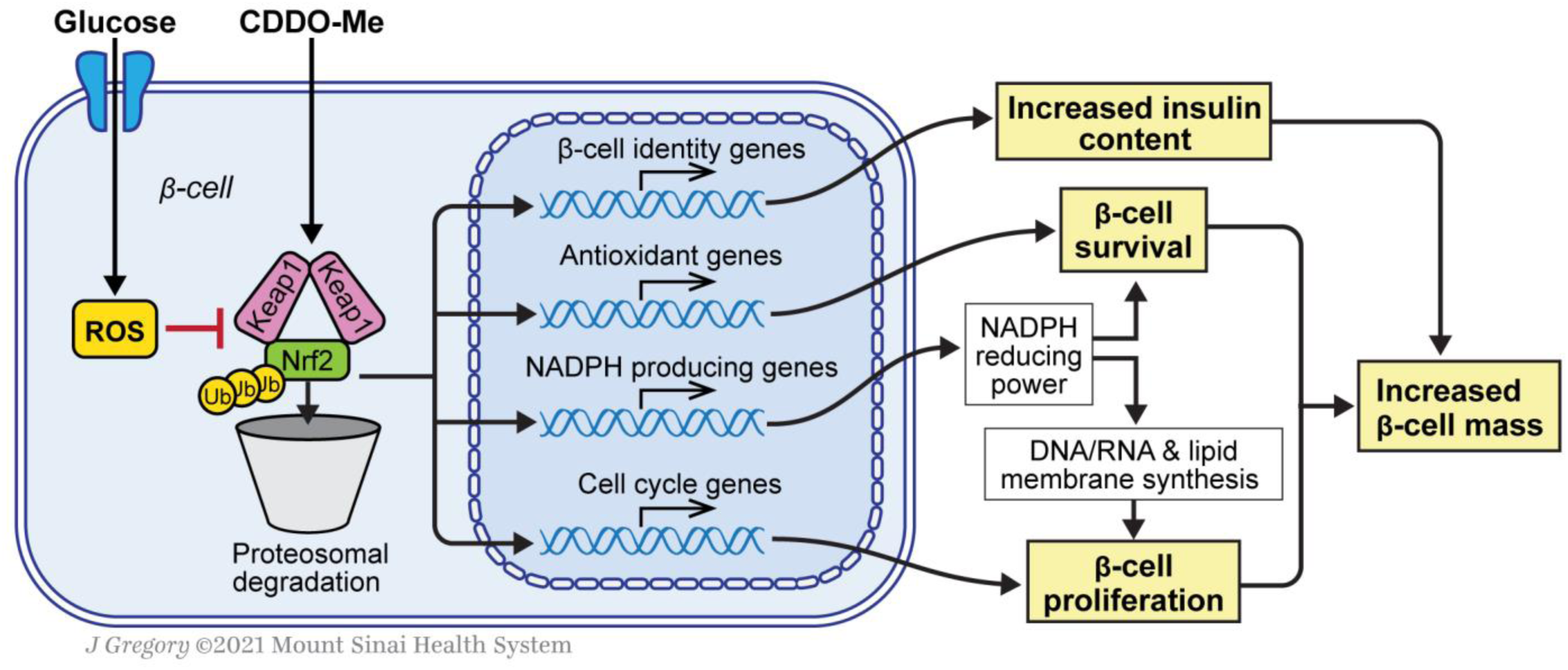
Nrf2 increases β-cell mass by increasing survival and promoting β-cell proliferation. Glucose metabolism generates ROS in β-cells, which changes Keap1 conformation, decreases Nrf2:Keap1 interaction and proteolytic degradation and increases Nrf2 abundance and transcriptional activity. CDDO-Me also activates Nrf2 by inhibiting Keap1. Nrf2 target genes include antioxidant enzymes and enzymes that produce NADPH, which is used by antioxidant enzymes to protect β-cells from ROS, and also as reducing equivalents for synthesis of RNA, DNA and membrane synthesis in proliferating cells. Activation of Nrf2 is also associated with increased expression of cell cycle regulators and increased insulin content. Thus, the combined actions of Nrf2, protection from apoptosis and increased proliferation, lead to increased β-cell mass.

These findings prompted us to test whether activation of Nrf2 in primary mouse and human β-cells by CDDO-Me would stimulate β-cell proliferation, as we have seen in β-cells with genetic deletion of Keap1. Like CDDO-Me, the broccoli-extracted isothiocyanate sulforaphane (SFN) is also an Nrf2 activator that targets Keap1 (Dinkova-Kostova et al., 2017a; Sova and Saso, 2018). In order to test the effect of CDDO-Me and SFN compounds on β-cell proliferation, mouse islets were isolated and incubated with increasing doses of CDDO-Me (0.2-20 nM) and SFN (0.2-20 nM) in media containing 5.5 mM glucose. Following 72 h, cells were fixed and immunostained for insulin and Ki67. Treatment with either 20 nM CDDO-Me or 20 nM SFN increased β-cell proliferation compared to vehicle control (7.1 and 6.3-fold increase, respectively) to levels comparable to high (20 mM) glucose (Fig 6A,B). Moreover, deletion of Nrf2 in Nrf2^lox/lox^ mouse islets, blocked CDDO-Me and SFN-stimulated β-cell proliferation (Fig 6C), indicating that these compounds stimulate β-cell proliferation by activation of Nrf2. Importantly, cadaveric human islets treated with either 20 nM CDDO-Me or 20 nM SFN also displayed increased β-cell proliferation (9.2 and 9.8-fold increase, respectively), similar to high glucose (Fig 6D,E). CDDO-Me and SFN at these concentrations did not induce human β-cell death as seen by TUNEL assay (Fig 6F).

We next tested the effect of a systemic *in vivo* administration of CDDO-Me on both mouse and human β-cell proliferation. For this purpose, euglycemic immunodeficient mice were transplanted with 500 human islets under the kidney capsule. Following 17 days of recovery, mice were IP injected daily with increasing doses of CDDO-Me for 7 days, after which mouse pancreata and the kidneys containing the grafts were harvested (Fig. 6G). Daily injection with 3 mg/kg CDDO-Me stimulated mouse β-cell proliferation (29.5-fold) (Fig. 6H,I). More importantly, daily injection with 1 mg/kg CDDO-Me significantly and markedly stimulated human β-cell proliferation (3.9-fold) compared to vehicle-treated transplanted mice (Fig. 6J,K,). Thus, pharmacological activation of Nrf2 leads to rodent and human β-cell proliferation *in vivo*.

## DISCUSSION

Nrf2 is a master regulator of transcriptional programs designed to protect cells from oxidative stress. Whereas it is well established that activation of the Nrf2 pathway protects β-cells from a variety of metabolic stresses (Baumel-Alterzon et al., 2020), here we add to these observations by demonstrating a pivotal role for Nrf2 in the regulation of β-cell mass. Our study reveals a number of novel findings including: 1) glucose rapidly activates Nrf2 in INS1 cells and primary β-cells, establishing Nrf2 as a glucose-regulated transcription factor in β-cells; 2) Nrf2 is necessary for glucose-stimulated β-cell proliferation and for adaptive β-cell proliferation, adaptive expansion of β-cell mass, and protection from cell death in mice on a HFD; 3) Nrf2 is necessary to maintain insulin content; 4) activation of Nrf2 is sufficient to increase β-cell proliferation in mouse and human β-cell *in vitro* and *in vivo*. Thus, our data support the notion that Nrf2 is a crucial regulator of β-cell mass (Figure 7).

We found that exposure of β-cells to high glucose results in increased Nrf2 nuclear localization and transcriptional activation. Thus, Nrf2 joins a growing list of glucose-responsive transcription factors in β-cells that increase transactivation capacity in response to increased glucose metabolism, including ChREBP, MondoA, and Myc (Abdul-Wahed et al., 2017; Rosselot et al., 2020; Zhang et al., 2010). ChREBP and MondoA are activated by binding metabolites of glycolysis, likely Glucose-6-phosphate (Abdul-Wahed et al., 2017), and Myc is stabilized by phosphorylation and phosphatase cascades initiated from glucose metabolism (Rosselot et al., 2020). Nrf2 is activated by increased production of ROS, which are generated mostly in the mitochondria as a byproduct of increased glucose metabolism (Baumel-Alterzon et al., 2020). Keap1:Nrf2 interaction is compromised by increased ROS via interaction with redox-sensitive cysteines (mainly cysteine C151), thus reducing Nrf2 ubiquitination and subsequent degradation (Dinkova-Kostova et al., 2017b; Fourquet et al., 2010; Panieri and Saso, 2019). It is possible that other glucose metabolites may activate Nrf2. For example, Bollong et al. recently found that methylglyoxal, a metabolite of glycolysis (Kim et al., 2004), forms covalent modifications on Keap1 cysteine C151, leading to dimerization and Nrf2 activation (Bollong et al., 2018). We cannot rule out methylglyoxal as a mediator of Nrf2 activation in β-cells, however, methylglyoxal itself can also generate ROS in β-cells (Bo et al., 2016) thus supporting the notion that glucose activates Nrf2 *via* production of ROS.

The Nrf2 protein contains three nuclear localization signals (NLS). The first is a monopartite NLS (NLS1) that is located at the N terminal region within the Neh2 domain. The second one is a bipartite NLS located at the Neh1 domain (NLS2), and a third one is a monopartite NLS that is located at the C terminal region (NLS3) (Giudice et al., 2010; Theodore et al., 2008). In addition, Nrf2 contains two nuclear export signals (NES). The first is a redox-insensitive NES located within the leucine zipper (ZIP) of the Neh1 domain (NES_ZIP_). The other is a redox-sensitive NES (NES_TA_) located in the Neh5 domain (Giudice et al., 2010; Li et al., 2006). During basal conditions, the combined activities of NES_ZIP_ and NES_TA_ counterbalance the NLS activity, leaving Nrf2 in the cytoplasm. However, under oxidative stress, cysteine C183 of NES_TA_ is modified, thus inhibiting the binding to nuclear export protein CRM1 (Li et al., 2006). The βNrf2KO mice that were used in this study utilized Cre-mediated deletion of Nrf2 exon 5, which encodes the Neh1 domain, necessary for Nrf2 binding to its obligate heterodimer partner, sMaf, and to DNA (Tonelli et al., 2018). Here we show that deletion of exon 5, which renders Nrf2 without NLS2 and NES_ZIP,_ but still contains NLS1,3 and NES_TA_, did not prevent Nrf2 translocation into the nucleus at high glucose. These observations are consistent with the idea that NES_TA_ is probably disabled at high glucose due to ROS generation.

Using a β-cell-specific conditional knockout model, we found that Nrf2 is necessary for glucose-stimulated β-cell proliferation *in vitro* and for adaptive proliferation and expansion of β-cell mass in response to a HFD *in vivo*. These results are consistent with and extend our previous results demonstrating that Nrf2 is activated in response to overexpression of ChREBPα, leading to increased anabolic metabolism, mitochondrial activity and glucose-stimulated β-cell proliferation (Kumar et al., 2018). As a master transcriptional regulator of redox balance, Nrf2 is uniquely positioned to support β-cell proliferation. Target genes of Nrf2 include antioxidant enzymes that protect β-cells from the increased oxidative stress brought about by increased mitochondrial activity and anabolic metabolism necessary for manufacturing the biomass needed to create daughter cells in the process of cell division (Baumel-Alterzon et al., 2020). It follows that we found an increase in the expression of antioxidant enzyme genes, and genes that produce NADPH, providing reducing power for the antioxidant enzymes, and for producing nucleotide and membrane phospholipid synthesis (Hosios and Vander Heiden, 2018; Mitsuishi et al., 2012; Vander Heiden et al., 2009) Thus, Nrf2 promotes adaptive proliferation by orchestrating programs of gene expression that support the anabolic workload of dividing cells as well as providing protection from the oxidative stress derived from a hypercaloric diet and cellular replication.

Chronic exposure of β-cells to high levels of ROS results in β-cell malfunction and apoptotic cell death (Gerber and Rutter, 2017; Kaneto et al., 2002; Kohnert et al., 2012; Krippeit-Drews et al., 1999; Lee et al., 2011; Matsuoka et al., 1997; Miki et al., 2018; Rebelato et al., 2010, 2011). Several researchers have found that activation of Nrf2 protects or reverses damage caused by oxidative stress in β-cells (Dinic et al., 2016; Fernandez-Millan et al., 2016; Li et al., 2018; Masuda et al., 2015; Moens et al., 2020; Schultheis et al., 2019; Wada et al., 2019; Yagishita et al., 2014). For example, Abebe et al. placed female fatty Zucker fatty rats on a high fat diet for 9 days, resulting in robust oxidative damage and the beginnings of β-cell failure and dysregulated glucose homeostasis. By returning the animals to a standard chow diet, the β-cells repaired themselves with evidence of a sharp upregulation and expression of Nrf2 and its antioxidant target genes (Abebe et al., 2017). Similarly, Yagashita et al. used a β-cell-specific iNOS2-expressing mouse model to create oxidative stress resulting in β-cell dysfunction and glucose intolerance. Activation of Nrf2 by deletion of Keap1 in a β-cell-specific manner reversed the phenotype (Yagishita et al., 2014). Furthermore, the same group found that a global Keap1+/-mouse crossed to db/db diabetic mice, or administration of CDDO-Im, an Nrf2 activator, attenuated diabetes in the db/db mice (Uruno et al., 2013). In concordance with these studies, we found that β-cell-specific depletion of Nrf2 in mice fed on a HFD resulted in increased oxidative stress and apoptosis of β-cells. Thus, Nrf2 defends β-cells from the oxidative damage caused by metabolic stress in a variety of model systems.

Interestingly, we found that depletion of Nrf2 in β-cells reduced insulin immunostaining and insulin content. These findings correlated with a decreased mRNA expression of Ins2 and Pdx1, and a decreased immunostaining of Pdx1 in pancreata from β-cell-specific Nrf2 knock out mice. Although the β-cell identity transcription factor Pdx1 is not known to be a direct Nrf2 target, its level and binding activity to the insulin promoter, and therefore subsequent insulin production, is impaired in hyperglycemic conditions, presumably as a result of oxidative stress and ROS generation (Glavas et al., 2019; Guo et al., 2013; Harmon et al., 1999; Reimer and Ahren, 2002; Robertson, 2006). Apart from Pdx1, Nrf2 deletion also contributed to reduced mRNA expression of other β-cell identity genes, such as MafA, Glut2 and Nkx6.1. Similar to Pdx1, these factors are highly sensitive to any changes in redox balance that can reduce their levels and activity (Guo et al., 2013; Yaribeygi et al., 2020) Thus, the contribution of Nrf2 activity towards building β-cell mass includes maintaining insulin content.

We found that increasing Nrf2 activity, either by conditionally deleting the Nrf2 inhibitor, Keap1, or with the Nrf2 activator CDDO-Me, was sufficient to increase rodent and human β-cell proliferation both *in vitro* and *in vivo* [(Kumar et al., 2018; Uruno et al., 2013) and this work]. In the Keap1 KO mouse model, we found that 1 month of Nrf2 activation significantly increased β-cell mass. Histomorphometry analysis revealed significant increases in the largest and smallest quartile of islets. Since the formation of new islets (neogenesis) from pancreatic progenitor cells results in small islets (Roostalu et al., 2020), we cannot exclude neogenesis as another mechanism by which Nrf2 promotes β-cell expansion. Lineage-tracing experiments are needed to determine this possibility.

We found that a weeklong treatment with CDDO-Me increased both rodent and human β-cell proliferation in the same model of human islet transplantation. This raises the exciting possibility of Nrf2 activators as therapeutic agents for expanding β-cell mass in diabetic patients. CDDO-Me is a synthetic oleanane triterpenoid and a member of the bardoxolone family of drugs, which have been investigated in several clinical trials, including chronic kidney disease in diabetic patients (Chertow et al., 2018; Chin et al., 2018; Pergola et al., 2011; Rizk et al., 2019; Rossing et al., 2019; Toto, 2018). Although some clinical trials were terminated early than expected due to cardiovascular safety concerns, a new phase 2 clinical trial with CDDO-Me has recently been started, excluding at-risk patients, to better define the safety and efficacy of CDDO-Me (Robledinos-Anton et al., 2019; Toto, 2018). Compounds of this drug family increase islet viability, preserve β-cell numbers, improve β-cell insulin content and insulin secretion, accelerate macro-autophagy, protect β-cells from oxidative stress and reduce secretion of proinflammatory cytokines (Li et al., 2015; Li et al., 2014; Masuda et al., 2015; Uruno et al., 2013; Yagishita et al., 2014), all of which can result in preservation of functional β-cell mass. Our results add β-cell proliferation as an additional property of Nrf2 activation by CDDO-Me and SFN. While this effect was examined in β-cells, it is also possible that proliferation in other islet cell types, or throughout the body, might be increased by CDDO-Me. Therefore, more studies are needed to ensure the safety of systemic Nrf2 activators and efforts should be made to design Nrf2 activators that are targeted specificity to β-cells for increasing β-cell mass in diabetic patients.

In summary, Nrf2 is necessary for maintaining β-cell redox balance and survival and is sufficient when activated to drive rodent and human β-cell proliferation. Thus, Nrf2 is a central regulator of β-cell mass, making this pathway a promising candidate for the development of future therapeutics for the treatment of diabetes.

## ACKNOWLEDGMENTS

Funding sources: NIH/NIDDK, DK114338 (DKS); DK020541, DK105015, and DK126450, AG-O and a Mindich Child Health and Development Institute Pilot and Feasibility Grant; Mindich Post-doc Pilot Award (SA-B). We thank the Developmental Studies Hybridoma Bank, at The University of Iowa, Department of Biology, Iowa City, for providing human insulin antibody. We thank the Icahn School of Medicine at Mount Sinai Microscopy Core and the Einstein/Sinai Diabetes Center Human Islet and Adenovirus Core.

## AUTHOR CONTRIBUTIONS

Conceptualization, SA-B and DKS; Investigation, SA-B, LSK, GB, YL, CJ-P; Writing – Original Draft, S-AB and DKS; Writing – Review & Editing, S-AB, DKS, AG-O, SB; Funding Acquisition, DKS, SA-B, AG-O; Resources, DKS, AG-O, SB; Supervision, DKS, AG-O. DKS is the guarantor of this work and, as such, had full access to all the data in the study and takes responsibility for the integrity of the data and the accuracy of the data analysis.

## RESEARCH DESIGN AND METHODS

### Cell culture

INS1-derived 832/13 rats insulinomas cell line (Hohmeier et al., 2000) were cultured in RPMI 1640 medium supplemented with 10% fetal bovine serum, 50 IU/mL penicillin, 50 mg/L streptomycin, 10 mM HEPES, 2 mM L-glutamine, 1 mM sodium pyruvate, and 50 µM β--mercaptoethanol. Cells were grown in a 37°C incubator under a humidified atmosphere containing 5% CO_2_.

### RT-PCR and quantitative real time PCR

Total RNA was extracted from isolated β cells using the RNeasy Micro kit (Qiagen #74004) and converted into double stranded complementary DNA (cDNA) using random hexamer primer (ThermoFisher Scientific #N8080127). The sequences of primers that were used for quantitative real time PCR are listed in supplementary Table 1.

### Mouse lines

βNrf2KO and βKeap1KO mice were generated by crossing MIP-CreER^TAM^ mice (Wicksteed et al., 2010) with Nrf2^lox/lox^ (Reddy et al., 2011) or Keap1^lox/lox^ (Blake et al., 2010) mice (respectively). Nrf2^lox/lox^ Keap1 ^lox/lox^ mice were generated by crossing Nrf2^lox/lox^ and Keap1 ^lox/lox^ mice. Cre expression was validated by PCR using the primers F’: CGCGGTCTGGCAGTAAAAACTATC and R’: CCCACCGTCAGTACGTGAGATATC. Nrf2 deletion was validated by PCR using the primers F’: TCTTAGGCACCATTTGGGAGAG and R’: TACAGCAGGCATACCATTGTGG, whereas Keap1 deletion was validated using the primers F’: CGAGGAAGCGTTTGCTTTAC and R’: GAGTCACCGTAAGCCTGGTC. βNrf2KO, βKeap1KO and MIP-CreER^TAM^ male mice were injected intraperitoneally for 5 consecutive days with 75 µg/g tamoxifen (Tam) (Sigma-Aldrich #T5648) dissolved in corn oil, as described in ref. (Rosselot et al., 2019). All studies were performed with the approval of and in accordance with guidelines established by the Icahn School of Medicine at Mount Sinai Institutional Animal Care and Use Committee.

### Adenoviruses, and reagents

Dispersed Nrf2^lox/lox^, Keap1 ^lox/lox^ and Nrf2^lox/lox^ Keap1 ^lox/lox^ mouse islets were transduced with adenoviral vectors encoding for Cre or for LacZ at an MOI of 150 in serum-free RPMI media for 2 hours at which time the culture media was replaced with serum-containing media (as previously described in (Kumar et al., 2018). Cells were treated with N-acetyl Cysteine (NAC) (Sigma # A9165), CDDO-Me (Sigma # SMB00376) Sulforaphane (SFN) (Sigma # S4441) or Brusatol (Ark Pharm AK128303).

### Human and mouse islets

Mouse islets were isolated after collagenase P (Sigma, #11213865001) injection through the pancreatic duct, followed by digestion and separation by density gradient using Histopaque®-1077 (Sigma # 10771), as previously reported (Roccisana et al., 2005). Human donor cadaveric islets were received from the Integrated Islet Distribution Program (IIDP) (https://iidp.coh.org/overview.aspx). Specific details of human islet donors are provided in Supplementary Table 2. Mouse and human islets were dispersed using trypsin and cultured overnight before immunofluorescent staining was carried out.

### Immunostaining

For β-cell proliferation, dispersed islets or paraffin-embedded pancreatic sections were immunostained with antibodies for mouse (Dako #A056401, or Genetex #GTX39371) or human insulin (Developmental Studies Hybridoma Bank, University of Iowa GN-ID4) and Ki67 (Invitrogen #MA5-14520). For assessing deletion of Nrf2 *in vitro*, cells were stained with Nrf2 (C-Term) antibody (Santa Cruz sc-722) or Nrf2-p (Abcam #ab76026). For assessing Nrf2 deletion *in vivo*, sections were stained with Nrf2 (C-Term) antibody (Cayman Chemicals #10214) or Nqo1 antibody (Abcam #ab2346). Keap1 levels were assessed by staining with (Cell Signaling #8047s). For assessing oxidative stress, sections were stained with antibody against 8OHdG Anti-DNA/RNA Damage antibody (Abcam #ab62623). For assessing Pdx1 levels, sections were stained with insulin (R&D #MAB1417) and Pdx1 (Abcam #AB47308). β-Cell mass and islet morphometry were measured as an average of three insulin stained mouse pancreatic sections using ImageJ software (National Institutes of Health) as described in (Alonso et al., 2007).

### TUNEL assay

TUNEL labeling was preformed according to manufacture, using the DeadEnd Fluormetric TUNEL System (Cat#G3250, Promega)

### Trypan blue viability assay

Following treatment, INS1 cells were stained with negative stain Trypan blue and live cells were counted under a light microscope.

### Acute and chronic HFD Feeding

2-4 months old βNrf2KO and βKeap1KO Tam- or vehicle corn oil-injected mice were placed for 1 week or 1 month on a lard-based HFD (41% kcal from fat) (TD 96001; Harlan Teklad) or a regular diet (RD) (13.1% kcal from fat) (Purina PicoLab 5053; LabDiet) as described in (Lakshmipathi et al., 2016). At the end of each experiment, mice were monitored for their body weights, non-fasting blood glucose, and plasma insulin and their pancreas was harvested and processed for histological studies or islet isolation.

### Glucose homeostasis and insulin content

Blood glucose was determined by glucometer and plasma insulin by ELISA assay (Mercodia, #10-1249-01). Intraperitoneal glucose tolerance test (IpGTT) was performed on 16–18 h fasted mice injected intraperitoneally with 2 g glucose/kg, as described in (Lakshmipathi et al., 2016). Insulin tolerance test (ITT) was performed in random-fed mice injected intraperitoneally with human insulin (1.5 units/kg), as previously described (Lakshmipathi et al., 2016). GSIS was performed as previously described (Wang et al., 2015a). In brief, mouse islets were preincubated in Krebs-Ringer bicarbonate buffer supplemented with 10 mmol/l HEPES, 1% BSA, and 2.8 mmol/l glucose for 1 h at 37°C in a 5% CO2 incubator. After washing once with the same solution, islets were incubated in fresh Krebs-Ringer bicarbonate buffer supplemented with 1% BSA and either 2.8 or 16.7 mmol/l glucose for 30 min. Buffer was removed and frozen at −20°C for insulin measurement by insulin ELISA assay (Mercodia, #10-1249-01). For insulin content, Islets were then digested for 2 h in NaOH at 37°C, follow by neutralization with HCl. Insulin values were normalized to DNA content.

### Euglycemic human islet transplantation model

500 human islet equivalents (IEQ) from five different human cadaveric donors were transplanted under the renal capsule of five euglycemic 3 to 7 month old NOD-SCID or Rag1-/-immunodeficient mice as detailed previously (Wang et al., 2015a). Animals were allowed to recover for seventeen days, and then were randomly selected to be given daily intraperitoneally injections of 0.1 or 1.0 or 3.0 mg/kg CDDO-Me or vehicle (DMSO) for seven days. Animals were then sacrificed, kidneys and pancreas harvested, fixed, embedded, sectioned and immunostained for insulin and Ki67.

### Statistical Analysis

Data is presented in this study as means ±SE. Statistical analysis was performed using unpaired two-tailed t test, one-way ANOVA and Two-way ANOVA with Tukey’s multiple comparison test.

## REFERENCES

Abdul-Wahed, A., Guilmeau, S., and Postic, C. (2017). Sweet Sixteenth for ChREBP: Established Roles and Future Goals. Cell Metab 26, 324–341.

Abebe, T., Mahadevan, J., Bogachus, L., Hahn, S., Black, M., Oseid, E., Urano, F., Cirulli, V., and Robertson, R.P. (2017). Nrf2/antioxidant pathway mediates beta cell self-repair after damage by high-fat diet-induced oxidative stress. JCI insight 2.

Aguayo-Mazzucato, C., and Bonner-Weir, S. (2018). Pancreatic beta Cell Regeneration as a Possible Therapy for Diabetes. Cell Metab 27, 57–67.

Alonso, L.C., Yokoe, T., Zhang, P., Scott, D.K., Kim, S.K., O’Donnell, C.P., and Garcia-Ocana, A. (2007). Glucose infusion in mice: a new model to induce beta-cell replication. Diabetes 56, 1792–1801.

Assmann, A., Ueki, K., Winnay, J.N., Kadowaki, T., and Kulkarni, R.N. (2009). Glucose effects on beta-cell growth and survival require activation of insulin receptors and insulin receptor substrate 2. Mol Cell Biol 29, 3219–3228.

Baumel-Alterzon, S., Katz, L.S., Brill, G., Garcia-Ocana, A., and Scott, D.K. (2020). Nrf2: The Master and Captain of Beta Cell Fate. Trends Endocrinol Metab.

Blake, D.J., Singh, A., Kombairaju, P., Malhotra, D., Mariani, T.J., Tuder, R.M., Gabrielson, E., and Biswal, S. (2010). Deletion of Keap1 in the lung attenuates acute cigarette smoke-induced oxidative stress and inflammation. Am J Respir Cell Mol Biol 42, 524–536.

Bloom, D.A., and Jaiswal, A.K. (2003). Phosphorylation of Nrf2 at Ser40 by protein kinase C in response to antioxidants leads to the release of Nrf2 from INrf2, but is not required for Nrf2 stabilization/accumulation in the nucleus and transcriptional activation of antioxidant response element-mediated NAD(P)H:quinone oxidoreductase-1 gene expression. J Biol Chem 278, 44675–44682.

Bo, J., Xie, S., Guo, Y., Zhang, C., Guan, Y., Li, C., Lu, J., and Meng, Q.H. (2016). Methylglyoxal Impairs Insulin Secretion of Pancreatic beta-Cells through Increased Production of ROS and Mitochondrial Dysfunction Mediated by Upregulation of UCP2 and MAPKs. J Diabetes Res 2016, 2029854.

Bollong, M.J., Lee, G., Coukos, J.S., Yun, H., Zambaldo, C., Chang, J.W., Chin, E.N., Ahmad, I., Chatterjee, A.K., Lairson, L.L., et al. (2018). A metabolite-derived protein modification integrates glycolysis with KEAP1-NRF2 signalling. Nature 562, 600–604.

Bryan, H.K., Olayanju, A., Goldring, C.E., and Park, B.K. (2013). The Nrf2 cell defence pathway: Keap1-dependent and -independent mechanisms of regulation. Biochemical pharmacology 85, 705–717.

Chertow, G.M., Appel, G.B., Block, G.A., Chin, M.P., Coyne, D.W., Goldsberry, A., Kalantar-Zadeh, K., Meyer, C.J., Molitch, M.E., Pergola, P.E., et al. (2018). Effects of bardoxolone methyl on body weight, waist circumference and glycemic control in obese patients with type 2 diabetes mellitus and stage 4 chronic kidney disease. J Diabetes Complications 32, 1113–1117.

Chin, M.P., Bakris, G.L., Block, G.A., Chertow, G.M., Goldsberry, A., Inker, L.A., Heerspink, H.J.L., O’Grady, M., Pergola, P.E., Wanner, C., et al. (2018). Bardoxolone Methyl Improves Kidney Function in Patients with Chronic Kidney Disease Stage 4 and Type 2 Diabetes: Post-Hoc Analyses from Bardoxolone Methyl Evaluation in Patients with Chronic Kidney Disease and Type 2 Diabetes Study. Am J Nephrol 47, 40–47.

Chung, Y.K., Chi-Hung Or, R., Lu, C.H., Ouyang, W.T., Yang, S.Y., and Chang, C.C. (2015). Sulforaphane down-regulates SKP2 to stabilize p27(KIP1) for inducing antiproliferation in human colon adenocarcinoma cells. J Biosci Bioeng 119, 35–42.

Cleasby, A., Yon, J., Day, P.J., Richardson, C., Tickle, I.J., Williams, P.A., Callahan, J.F., Carr, R., Concha, N., Kerns, J.K., et al. (2014). Structure of the BTB domain of Keap1 and its interaction with the triterpenoid antagonist CDDO. PLoS One 9, e98896.

Crean, D., Felice, L., Taylor, C.T., Rabb, H., Jennings, P., and Leonard, M.O. (2012). Glucose reintroduction triggers the activation of Nrf2 during experimental ischemia reperfusion. Mol Cell Biochem 366, 231–238.

Dinic, S., Grdovic, N., Uskokovic, A., Dordevic, M., Mihailovic, M., Jovanovic, J.A., Poznanovic, G., and Vidakovic, M. (2016). CXCL12 protects pancreatic beta-cells from oxidative stress by a Nrf2-induced increase in catalase expression and activity. Proc Jpn Acad Ser B Phys Biol Sci 92, 436–454.

Dinkova-Kostova, A.T., Fahey, J.W., Kostov, R.V., and Kensler, T.W. (2017a). KEAP1 and Done? Targeting the NRF2 Pathway with Sulforaphane. Trends Food Sci Technol 69, 257–269.

Dinkova-Kostova, A.T., Kostov, R.V., and Canning, P. (2017b). Keap1, the cysteine-based mammalian intracellular sensor for electrophiles and oxidants. Arch Biochem Biophys 617, 84–93.

Fernandez-Millan, E., Martin, M.A., Goya, L., Lizarraga-Mollinedo, E., Escriva, F., Ramos, S., and Alvarez, C. (2016). Glucagon-like peptide-1 improves beta-cell antioxidant capacity via extracellular regulated kinases pathway and Nrf2 translocation. Free Radic Biol Med 95, 16–26.

Fourquet, S., Guerois, R., Biard, D., and Toledano, M.B. (2010). Activation of NRF2 by nitrosative agents and H2O2 involves KEAP1 disulfide formation. J Biol Chem 285, 8463–8471.

Fu, J., Xiong, Z., Huang, C., Li, J., Yang, W., Han, Y., Paiboonrungruan, C., Major, M.B., Chen, K.N., Kang, X., et al. (2019). Hyperactivity of the transcription factor Nrf2 causes metabolic reprogramming in mouse esophagus. J Biol Chem 294, 327–340.

Gerber, P.A., and Rutter, G.A. (2017). The Role of Oxidative Stress and Hypoxia in Pancreatic Beta-Cell Dysfunction in Diabetes Mellitus. Antioxid Redox Signal 26, 501–518.

Giudice, A., Arra, C., and Turco, M.C. (2010). Review of molecular mechanisms involved in the activation of the Nrf2-ARE signaling pathway by chemopreventive agents. Methods Mol Biol 647, 37–74.

Glavas, M.M., Hui, Q., Tuduri, E., Erener, S., Kasteel, N.L., Johnson, J.D., and Kieffer, T.J. (2019). Early overnutrition reduces Pdx1 expression and induces beta cell failure in Swiss Webster mice. Sci Rep 9, 3619.

Gracia-ocana, A., Alonso, L.A. (2010). Glucose Mediated Regulation of Beta Cell Proliferation. Open Endocrinol J 4, 55–65.

Guo, S., Dai, C., Guo, M., Taylor, B., Harmon, J.S., Sander, M., Robertson, R.P., Powers, A.C., and Stein, R. (2013). Inactivation of specific beta cell transcription factors in type 2 diabetes. J Clin Invest 123, 3305–3316.

Harder, B., Tian, W., La Clair, J.J., Tan, A.C., Ooi, A., Chapman, E., and Zhang, D.D. (2017). Brusatol overcomes chemoresistance through inhibition of protein translation. Mol Carcinog 56, 1493–1500.

Harmon, J.S., Gleason, C.E., Tanaka, Y., Oseid, E.A., Hunter-Berger, K.K., and Robertson, R.P. (1999). In vivo prevention of hyperglycemia also prevents glucotoxic effects on PDX-1 and insulin gene expression. Diabetes 48, 1995–2000.

Hayes, J.D., and Ashford, M.L. (2012). Nrf2 orchestrates fuel partitioning for cell proliferation. Cell Metab 16, 139–141.

Hohmeier, H.E., Mulder, H., Chen, G., Henkel-Rieger, R., Prentki, M., and Newgard, C.B. (2000). Isolation of INS-1-derived cell lines with robust ATP-sensitive K+ channel-dependent and -independent glucose-stimulated insulin secretion. Diabetes 49, 424–430.

Hosios, A.M., and Vander Heiden, M.G. (2018). The redox requirements of proliferating mammalian cells. J Biol Chem 293, 7490–7498.

Jiang, T., Huang, Z., Lin, Y., Zhang, Z., Fang, D., and Zhang, D.D. (2010). The protective role of Nrf2 in streptozotocin-induced diabetic nephropathy. Diabetes 59, 850–860.

Jimenez-Osorio, A.S., Gonzalez-Reyes, S., Garcia-Nino, W.R., Moreno-Macias, H., Rodriguez-Arellano, M.E., Vargas-Alarcon, G., Zuniga, J., Barquera, R., and Pedraza-Chaverri, J. (2016). Association of Nuclear Factor-Erythroid 2-Related Factor 2, Thioredoxin Interacting Protein, and Heme Oxygenase-1 Gene Polymorphisms with Diabetes and Obesity in Mexican Patients. Oxid Med Cell Longev 2016, 7367641.

Kaneto, H., Xu, G., Fujii, N., Kim, S., Bonner-Weir, S., and Weir, G.C. (2002). Involvement of c-Jun N-terminal kinase in oxidative stress-mediated suppression of insulin gene expression. J Biol Chem 277, 30010–30018.

Kim, J., Son, J.W., Lee, J.A., Oh, Y.S., and Shinn, S.H. (2004). Methylglyoxal induces apoptosis mediated by reactive oxygen species in bovine retinal pericytes. J Korean Med Sci 19, 95–100.

Kohnert, K.D., Freyse, E.J., and Salzsieder, E. (2012). Glycaemic variability and pancreatic beta-cell dysfunction. Curr Diabetes Rev 8, 345–354.

Komatsu, M., Kurokawa, H., Waguri, S., Taguchi, K., Kobayashi, A., Ichimura, Y., Sou, Y.S., Ueno, I., Sakamoto, A., Tong, K.I., et al. (2010). The selective autophagy substrate p62 activates the stress responsive transcription factor Nrf2 through inactivation of Keap1. Nature cell biology 12, 213–223.

Krippeit-Drews, P., Kramer, C., Welker, S., Lang, F., Ammon, H.P., and Drews, G. (1999). Interference of H2O2 with stimulus-secretion coupling in mouse pancreatic beta-cells. J Physiol 514 (Pt 2), 471–481.

Kumar, A., Katz, L.S., Schulz, A.M., Kim, M., Honig, L.B., Li, L., Davenport, B., Homann, D., Garcia-Ocana, A., Herman, M.A., et al. (2018). Activation of Nrf2 Is Required for Normal and ChREBPalpha-Augmented Glucose-Stimulated beta-Cell Proliferation. Diabetes 67, 1561–1575.

Lakshmipathi, J., Alvarez-Perez, J.C., Rosselot, C., Casinelli, G.P., Stamateris, R.E., Rausell-Palamos, F., O’Donnell, C.P., Vasavada, R.C., Scott, D.K., Alonso, L.C., et al. (2016). PKCzeta Is Essential for Pancreatic beta-Cell Replication During Insulin Resistance by Regulating mTOR and Cyclin-D2. Diabetes 65, 1283–1296.

Lee, E., Ryu, G.R., Ko, S.H., Ahn, Y.B., Yoon, K.H., Ha, H., and Song, K.H. (2011). Antioxidant treatment may protect pancreatic beta cells through the attenuation of islet fibrosis in an animal model of type 2 diabetes. Biochem Biophys Res Commun 414, 397–402.

Li, C.G., Ni, C.L., Yang, M., Tang, Y.Z., Li, Z., Zhu, Y.J., Jiang, Z.H., Sun, B., and Li, C.J. (2018). Honokiol protects pancreatic beta cell against high glucose and intermittent hypoxia-induced injury by activating Nrf2/ARE pathway in vitro and in vivo. Biomed Pharmacother 97, 1229–1237.

Li, J., Zhang, C., Xing, Y., Janicki, J.S., Yamamoto, M., Wang, X.L., Tang, D.Q., and Cui, T. (2011). Up-regulation of p27(kip1) contributes to Nrf2-mediated protection against angiotensin II-induced cardiac hypertrophy. Cardiovasc Res 90, 315–324.

Li, M., Yu, H., Pan, H., Zhou, X., Ruan, Q., Kong, D., Chu, Z., Li, H., Huang, J., Huang, X., et al. (2019). Nrf2 Suppression Delays Diabetic Wound Healing Through Sustained Oxidative Stress and Inflammation. Front Pharmacol 10, 1099.

Li, S., Vaziri, N.D., Masuda, Y., Hajighasemi-Ossareh, M., Robles, L., Le, A., Vo, K., Chan, J.Y., Foster, C.E., Stamos, M.J., et al. (2015). Pharmacological activation of Nrf2 pathway improves pancreatic islet isolation and transplantation. Cell Transplant 24, 2273–2283.

Li, W., Wu, W., Song, H., Wang, F., Li, H., Chen, L., Lai, Y., Janicki, J.S., Ward, K.W., Meyer, C.J., et al. (2014). Targeting Nrf2 by dihydro-CDDO-trifluoroethyl amide enhances autophagic clearance and viability of beta-cells in a setting of oxidative stress. FEBS letters 588, 2115–2124.

Li, W., Yu, S.W., and Kong, A.N. (2006). Nrf2 possesses a redox-sensitive nuclear exporting signal in the Neh5 transactivation domain. J Biol Chem 281, 27251–27263.

Lim, S., and Kaldis, P. (2013). Cdks, cyclins and CKIs: roles beyond cell cycle regulation. Development 140, 3079–3093.

Lo, S.C., and Hannink, M. (2008). PGAM5 tethers a ternary complex containing Keap1 and Nrf2 to mitochondria. Exp Cell Res 314, 1789–1803.

Malhotra, D., Portales-Casamar, E., Singh, A., Srivastava, S., Arenillas, D., Happel, C., Shyr, C., Wakabayashi, N., Kensler, T.W., Wasserman, W.W., et al. (2010). Global mapping of binding sites for Nrf2 identifies novel targets in cell survival response through ChIP-Seq profiling and network analysis. Nucleic Acids Res 38, 5718–5734.

Masuda, Y., Vaziri, N.D., Li, S., Le, A., Hajighasemi-Ossareh, M., Robles, L., Foster, C.E., Stamos, M.J., Al-Abodullah, I., Ricordi, C., et al. (2015). The effect of Nrf2 pathway activation on human pancreatic islet cells. PLoS One 10, e0131012.

Matana, A., Ziros, P.G., Chartoumpekis, D.V., Renaud, C.O., Polasek, O., Hayward, C., Zemunik, T., and Sykiotis, G.P. (2020). Rare and common genetic variations in the Keap1/Nrf2 antioxidant response pathway impact thyroglobulin gene expression and circulating levels, respectively. Biochem Pharmacol 173, 113605.

Matsuoka, T., Kajimoto, Y., Watada, H., Kaneto, H., Kishimoto, M., Umayahara, Y., Fujitani, Y., Kamada, T., Kawamori, R., and Yamasaki, Y. (1997). Glycation-dependent, reactive oxygen species-mediated suppression of the insulin gene promoter activity in HIT cells. J Clin Invest 99, 144–150.

Mazumder, S., DuPree, E.L., and Almasan, A. (2004). A dual role of cyclin E in cell proliferation and apoptosis may provide a target for cancer therapy. Curr Cancer Drug Targets 4, 65–75.

Meier, J.J., Butler, A.E., Saisho, Y., Monchamp, T., Galasso, R., Bhushan, A., Rizza, R.A., and Butler, P.C. (2008). Beta-cell replication is the primary mechanism subserving the postnatal expansion of beta-cell mass in humans. Diabetes 57, 1584–1594.

Miki, A., Ricordi, C., Sakuma, Y., Yamamoto, T., Misawa, R., Mita, A., Molano, R.D., Vaziri, N.D., Pileggi, A., and Ichii, H. (2018). Divergent antioxidant capacity of human islet cell subsets: A potential cause of beta-cell vulnerability in diabetes and islet transplantation. PLoS One 13, e0196570.

Mitsuishi, Y., Taguchi, K., Kawatani, Y., Shibata, T., Nukiwa, T., Aburatani, H., Yamamoto, M., and Motohashi, H. (2012). Nrf2 redirects glucose and glutamine into anabolic pathways in metabolic reprogramming. Cancer Cell 22, 66–79.

Moens, C., Bensellam, M., Himpe, E., Muller, C.J.F., Jonas, J.C., and Bouwens, L. (2020). Aspalathin Protects Insulin-Producing beta Cells against Glucotoxicity and Oxidative Stress-Induced Cell Death. Mol Nutr Food Res, e1901009.

Mosser, R.E., Maulis, M.F., Moulle, V.S., Dunn, J.C., Carboneau, B.A., Arasi, K., Pappan, K., Poitout, V., and Gannon, M. (2015). High-fat diet-induced beta-cell proliferation occurs prior to insulin resistance in C57Bl/6J male mice. American journal of physiology. Endocrinology and metabolism 308, E573–582.

Moulle, V.S., Vivot, K., Tremblay, C., Zarrouki, B., Ghislain, J., and Poitout, V. (2017). Glucose and fatty acids synergistically and reversibly promote beta cell proliferation in rats. Diabetologia 60, 879–888.

Mullarky, E., and Cantley, L.C. (2015). Diverting Glycolysis to Combat Oxidative Stress. In Innovative Medicine: Basic Research and Development. K. Nakao, N. Minato, and S. Uemoto, eds. (Tokyo), pp. 3–23.

Newsholme, P., Keane, K.N., Carlessi, R., and Cruzat, V. (2019). Oxidative stress pathways in pancreatic beta-cells and insulin-sensitive cells and tissues: importance to cell metabolism, function, and dysfunction. Am J Physiol Cell Physiol 317, C420–C433.

Panieri, E., and Saso, L. (2019). Potential Applications of NRF2 Inhibitors in Cancer Therapy. Oxid Med Cell Longev 2019, 8592348.

Pergola, P.E., Raskin, P., Toto, R.D., Meyer, C.J., Huff, J.W., Grossman, E.B., Krauth, M., Ruiz, S., Audhya, P., Christ-Schmidt, H., et al. (2011). Bardoxolone methyl and kidney function in CKD with type 2 diabetes. N Engl J Med 365, 327–336.

Pils, D., Bachmayr-Heyda, A., Auer, K., Svoboda, M., Auner, V., Hager, G., Obermayr, E., Reiner, A., Reinthaller, A., Speiser, P., et al. (2014). Cyclin E1 (CCNE1) as independent positive prognostic factor in advanced stage serous ovarian cancer patients - a study of the OVCAD consortium. Eur J Cancer 50, 99–110.

Rebelato, E., Abdulkader, F., Curi, R., and Carpinelli, A.R. (2010). Low doses of hydrogen peroxide impair glucose-stimulated insulin secretion via inhibition of glucose metabolism and intracellular calcium oscillations. Metabolism 59, 409–413.

Rebelato, E., Abdulkader, F., Curi, R., and Carpinelli, A.R. (2011). Control of the intracellular redox state by glucose participates in the insulin secretion mechanism. PLoS One 6, e24507.

Reddy, N.M., Potteti, H.R., Mariani, T.J., Biswal, S., and Reddy, S.P. (2011). Conditional deletion of Nrf2 in airway epithelium exacerbates acute lung injury and impairs the resolution of inflammation. American journal of respiratory cell and molecular biology 45, 1161–1168.

Reimer, M.K., and Ahren, B. (2002). Altered beta-cell distribution of pdx-1 and GLUT-2 after a short-term challenge with a high-fat diet in C57BL/6J mice. Diabetes 51 *Suppl 1*, S138–143.

Rizk, D.V., Silva, A.L., Pergola, P.E., Toto, R., Warnock, D.G., Chin, M.P., Goldsberry, A., O’Grady, M., Meyer, C.J., and McCullough, P.A. (2019). Effects of Bardoxolone Methyl on Magnesium in Patients with Type 2 Diabetes Mellitus and Chronic Kidney Disease. Cardiorenal Med 9, 316–325.

Robertson, R.P. (2006). Oxidative stress and impaired insulin secretion in type 2 diabetes. Curr Opin Pharmacol 6, 615–619.

Robledinos-Anton, N., Fernandez-Gines, R., Manda, G., and Cuadrado, A. (2019). Activators and Inhibitors of NRF2: A Review of Their Potential for Clinical Development. Oxid Med Cell Longev 2019, 9372182.

Roccisana, J., Reddy, V., Vasavada, R.C., Gonzalez-Pertusa, J.A., Magnuson, M.A., and Garcia-Ocana, A. (2005). Targeted inactivation of hepatocyte growth factor receptor c-met in beta-cells leads to defective insulin secretion and GLUT-2 downregulation without alteration of beta-cell mass. Diabetes 54, 2090–2102.

Rojas, J., Bermudez, V., Palmar, J., Martinez, M.S., Olivar, L.C., Nava, M., Tomey, D., Rojas, M., Salazar, J., Garicano, C., et al. (2018). Pancreatic Beta Cell Death: Novel Potential Mechanisms in Diabetes Therapy. J Diabetes Res 2018, 9601801.

Roostalu, U., Lercke Skytte, J., Gravesen Salinas, C., Klein, T., Vrang, N., Jelsing, J., and Hecksher-Sorensen, J. (2020). 3D quantification of changes in pancreatic islets in mouse models of diabetes type I and II. Dis Model Mech 13.

Rosselot, C., Baumel-Alterzon, S., Li, Y., Brill, G., Lambertini, L., Katz, L.S., Lu, G., Garcia-Ocana, A., and Scott, D.K. (2020). The Many Lives of Myc in the Pancreatic beta-Cell. J Biol Chem.

Rosselot, C., Kumar, A., Lakshmipathi, J., Zhang, P., Lu, G., Katz, L.S., Prochownik, E.V., Stewart, A.F., Lambertini, L., Scott, D.K., et al. (2019). Myc Is Required for Adaptive beta-Cell Replication in Young Mice but Is Not Sufficient in One-Year-Old Mice Fed With a High-Fat Diet. Diabetes 68, 1934–1949.

Rossing, P., Block, G.A., Chin, M.P., Goldsberry, A., Heerspink, H.J.L., McCullough, P.A., Meyer, C.J., Packham, D., Pergola, P.E., Spinowitz, B., et al. (2019). Effect of bardoxolone methyl on the urine albumin-to-creatinine ratio in patients with type 2 diabetes and stage 4 chronic kidney disease. Kidney Int 96, 1030–1036.

Schultheis, J., Beckmann, D., Mulac, D., Muller, L., Esselen, M., and Dufer, M. (2019). Nrf2 Activation Protects Mouse Beta Cells from Glucolipotoxicity by Restoring Mitochondrial Function and Physiological Redox Balance. Oxid Med Cell Longev 2019, 7518510.

Sova, M., and Saso, L. (2018). Design and development of Nrf2 modulators for cancer chemoprevention and therapy: a review. Drug Des Devel Ther 12, 3181–3197.

Stamateris, R.E., Sharma, R.B., Hollern, D.A., and Alonso, L.C. (2013). Adaptive beta-cell proliferation increases early in high-fat feeding in mice, concurrent with metabolic changes, with induction of islet cyclin D2 expression. American journal of physiology. Endocrinology and metabolism 305, E149–159.

Stamateris, R.E., Sharma, R.B., Kong, Y., Ebrahimpour, P., Panday, D., Ranganath, P., Zou, B., Levitt, H., Parambil, N.A., O’Donnell, C.P., et al. (2016). Glucose Induces Mouse beta-Cell Proliferation via IRS2, MTOR, and Cyclin D2 but Not the Insulin Receptor. Diabetes 65, 981–995.

Suzuki, T., and Yamamoto, M. (2017). Stress-sensing mechanisms and the physiological roles of the Keap1-Nrf2 system during cellular stress. J Biol Chem 292, 16817–16824.

Tamberg, N., Tahk, S., Koit, S., Kristjuhan, K., Kasvandik, S., Kristjuhan, A., and Ilves, I. (2018). Keap1-MCM3 interaction is a potential coordinator of molecular machineries of antioxidant response and genomic DNA replication in metazoa. Sci Rep 8, 12136.

Theodore, M., Kawai, Y., Yang, J., Kleshchenko, Y., Reddy, S.P., Villalta, F., and Arinze, I.J. (2008). Multiple nuclear localization signals function in the nuclear import of the transcription factor Nrf2. J Biol Chem 283, 8984–8994.

Tonelli, C., Chio, I.I.C., and Tuveson, D.A. (2018). Transcriptional Regulation by Nrf2. Antioxidants & redox signaling 29, 1727–1745.

Toto, R.D. (2018). Bardoxolone-the Phoenix? J Am Soc Nephrol 29, 360–361.

Uruno, A., Furusawa, Y., Yagishita, Y., Fukutomi, T., Muramatsu, H., Negishi, T., Sugawara, A., Kensler, T.W., and Yamamoto, M. (2013). The Keap1-Nrf2 system prevents onset of diabetes mellitus. Mol Cell Biol 33, 2996–3010.

Valavanidis, A., Vlachogianni, T., and Fiotakis, C. (2009). 8-hydroxy-2’ -deoxyguanosine (8-OHdG): A critical biomarker of oxidative stress and carcinogenesis. J Environ Sci Health C Environ Carcinog Ecotoxicol Rev 27, 120–139.

Vander Heiden, M.G., Cantley, L.C., and Thompson, C.B. (2009). Understanding the Warburg effect: the metabolic requirements of cell proliferation. Science 324, 1029–1033.

Wada, Y., Takata, A., Ikemoto, T., Morine, Y., Imura, S., Iwahashi, S., Saito, Y., and Shimada, M. (2019). The protective effect of epigallocatechin 3-gallate on mouse pancreatic islets via the Nrf2 pathway. Surg Today 49, 536–545.

Wan, Z.H., Jiang, T.Y., Shi, Y.Y., Pan, Y.F., Lin, Y.K., Ma, Y.H., Yang, C., Feng, X.F., Huang, L.F., Kong, X.N., et al. (2020). RPB5-Mediating Protein Promotes Cholangiocarcinoma Tumorigenesis and Drug Resistance by Competing With NRF2 for KEAP1 Binding. Hepatology 71, 2005–2022.

Wang, J., and Wang, H. (2017). Oxidative Stress in Pancreatic Beta Cell Regeneration. Oxid Med Cell Longev 2017, 1930261.

Wang, P., Alvarez-Perez, J.C., Felsenfeld, D.P., Liu, H., Sivendran, S., Bender, A., Kumar, A., Sanchez, R., Scott, D.K., Garcia-Ocana, A., et al. (2015a). A high-throughput chemical screen reveals that harmine-mediated inhibition of DYRK1A increases human pancreatic beta cell replication. Nature medicine 21, 383–388.

Wang, X., Chen, H., Liu, J., Ouyang, Y., Wang, D., Bao, W., and Liu, L. (2015b). Association between the NF-E2 Related Factor 2 Gene Polymorphism and Oxidative Stress, Anti-Oxidative Status, and Newly-Diagnosed Type 2 Diabetes Mellitus in a Chinese Population. Int J Mol Sci 16, 16483–16496.

Wicksteed, B., Brissova, M., Yan, W., Opland, D.M., Plank, J.L., Reinert, R.B., Dickson, L.M., Tamarina, N.A., Philipson, L.H., Shostak, A., et al. (2010). Conditional gene targeting in mouse pancreatic ss-Cells: analysis of ectopic Cre transgene expression in the brain. Diabetes 59, 3090–3098.

Yagishita, Y., Fukutomi, T., Sugawara, A., Kawamura, H., Takahashi, T., Pi, J., Uruno, A., and Yamamoto, M. (2014). Nrf2 protects pancreatic beta-cells from oxidative and nitrosative stress in diabetic model mice. Diabetes 63, 605–618.

Yaribeygi, H., Sathyapalan, T., Atkin, S.L., and Sahebkar, A. (2020). Molecular Mechanisms Linking Oxidative Stress and Diabetes Mellitus. Oxid Med Cell Longev 2020, 8609213.

Zhang, P., Metukuri, M.R., Bindom, S.M., Prochownik, E.V., O’Doherty, R.M., and Scott, D.K. (2010). c-Myc is required for the CHREBP-dependent activation of glucose-responsive genes. Mol Endocrinol 24, 1274–1286.

