## Supplemental Materials for "Nrf2 Regulates β-cell Mass by Suppressing Cell Death and Promoting Proliferation"

**Table 1: Primers sequences used in this paper for qPCR**

| Gene Name | Forward sequence | Reverse Sequence |
|---|---|---|
| ACTIN | AGCCATGTACGTAGCCATCC | CTCTCAGCTGTGGTGGTGAA |
| NRF2 | AGGACATGGAGCAAGTTTGG | TTCTTTTTCCAGCGAGGAGA |
| NQO1 | GAAGGAGGCTGCTGTAGAGG | ATCACCAGGTCTGCAGCTTC |
| KEAP1 | GATCGGCTGCACTGAACTG | GGACTCGCAGCGTACGTT |
| CCNA1 | CACAGAGAACCGTGCTAGGG | CACTTTCTTTCCAGCCGCAG |
| CCNA2 | GAGGGCGATCCTTGTGGAT | CACAGCCAAATGCAGGGTCT |
| CCNB1 | TGTGTGTGAACCAGAGGTGGAACT | AGATGTTTCCATCGGGCTTGGAGA |
| CCNB2 | GGCTGGTCCAAGTCCATTCC | GTCCATGATGGCAATGCACA |
| CCNB3 | TTGCTGAACTTCCGATCCCA | TTGCTTGCTCAAAGGAGGGA |
| CCND1 | CAGAAGTGCGAAGAGGAGGTC | TCATCTTAGAGGCCACGAACAT |
| CCND2 | TGTGGATTGTCTCAAAGCCTG | CAACATCCCGCACGTCTGTA |
| CCNE1 | GACACAGCTTCGGGGTGC | AACTCAGACCTGGGAGGACA |
| CCNE2 | TGCTGCCGCCTTATGTCATT | TCGGAGATGTCATCCCATTCC |
| CDK1 | CTGCAGCTCGGAGCACAGTT | CCAGAACACGGAGGCACTTG |
| CDK2 | AGCCAAGTTTCCCCAAGTGG | TTTGGGAAGGGCATCAGAGC |
| CDK4 | AGACCAGGACCTGAGGACAT | TCAGGTCCCGGTGAACAATG |
| CDK6 | GGCTATGGGAAGGTGTTCAA | GGGCTCTGGAACTTTATCCA |
| CDC25A | CACGGGGAAGATGCTGTTTG | AACAAGACAGGATGCCCAGC |
| P15 | GGGGGCAAGTGGAGACGG | CTTCCCGAGCTGCGTCGT |
| P16 | GTCTTTGTGTACCGCTGGGA | GCCGGATTTAGCTCTGCTCT |
| P18 | ACGTCAACGCTCAAAATGGA | TGGGATTAGCACCTCTGAGGA |
| P21 | GTGGCCTTGTCGCTGTCTT | GCGCTTGGAGTGATAGAAATCTG |
| P27 | GGTGGACCAAATGCCTGACT | TGGCCCTTTTGTTTTGCGAA |
| P57 | GGTGTCCCTCTCCAAACGTG | TGCCCAGCAAGTTCTCTCTG |
| INS1 | TATAAAGCTGGTGGGCATCC | GGGACCACAAAGATGCTGT |
| INS2 | TTTGTCAAGCAGCACCTTT | AGGTTTTCTCGCCCCTTAAC |
| MAFA | ATCATCACTCTGCCCACCAT | AGTCGGATGACCTCCTCCTT |
| GLUT2 | GGCACAGACACCCCACTTAC | GCCAACATTGCTTTGATCCT |
| PDX1 | CTCCGGACATCTCCCCATAC | ACGGGTCCTCTTGTTTTCCT |
| NKX6.1 | CGCCCGGGCTCTACTTTAG | GTCCAGAGAACGTGGGTCTG |
| PGD | CTCGGTGCTTGTCCTCTCTG | TTGAGGGTCCAGCCAAACTC |
| G6PD | CAACAGGATCTTTGGCCCCA | ACGGACATCATCTGAACCTGT |
| IDH1 | AGCTTCATCGCAACCCAGAA | GTAATAAGCCGGCCCCAGTT |

**Table 2: Human donor cadaveric islets used in this paper**

| Donor ID | Age (years) | Sex | Race | HbA1c (%) | BMI | COVID19 | Cause of Death | Used in |
|---|---|---|---|---|---|---|---|---|
| **SAMN1295912** | 37 | F | C | 5.3 | 26.3 | N/A | S | Fig. 7D-F |
| **SAMN13570019** | 37 | F | C | 5.2 | 24 | N/A | A | Fig. 7D-F |
| **SAMN13881228** | 34 | M | H | 5.6 | 28.1 | N/A | HT | Fig. 7D-F |
| **SMN12713942** | 41 | M | C | 5.2 | 23.5 | N/A | S | Fig. 7D-F |
| **SAMN12924398** | 38 | M | H | 5.4 | 28 | N/A | HT | Fig. 7D-F |
| **SAMN1295912** | 37 | F | C | 5.3 | 26.3 | N/A | S | Fig. 7D-F |
| **SAMN16427178** | 42 | F | C | N/A | 31.2 | Negative | S | Fig. 7J-K |
| **SAMN13881228** | 34 | M | H | N/A | 28.1 | N/A | HT | Fig. 7J-K |
| **HP-20220-01** | 48 | M | AF | 5.2 | 20.2 | N/A | S | Fig. 7J-K |
| **HP-20152-01** | 21 | M | H | 5.1 | 27.3 | Negative | HT | Fig. 7J-K |
| **HP-20199-01** | 48 | F | C | 5.8 | 30.9 | Negative | S | Fig. 7J-K |

M = males; F = female; H = Hispanic; C = Caucasian; AF = African American; HT = Head trauma; S = Stroke; A = Anoxia

**Supp. Figure 1. Acute exposure of INS1 to high glucose stimulates β-cell proliferation.** (A) INS-1 832/13 cells were cultured in 2 mM or 20 mM glucose for 5 min in the presence or absence of the antioxidant agent N-acetylcysteine (NAC) (20 mM). 100 μM $H_2O_2$ was used as a positive control. The cells were fixed and immunolabeled using an Nrf2-p antibody. (B) INS-1 832/13 cells were incubated in 2 mM or 20 mM glucose for 6 h. RNA was extracted and expression of Nqo1 was measured. (C,D) INS-1 832/13 cells were incubated in 2 mM or 20 mM glucose and with indicated concentrations of brusatol for 72 h. Cells were fixed and immunolabeled using Nrf2 or Ki67 antibodies. Data shown are the mean +/- SE (n=3-6, *p < 0.05; **p < 0.005; ****p < 0.0001).

**Supp. Figure 2. *In vivo* loss of Nrf2 function decreases HFD-stimulated β-cell proliferation.** (A) Mice were fed with HFD or RD. After one week, mice were euthanized and their pancreata were stained with insulin and Keap1 antibodies. Mean intensity was then calculated for Keap1. (B) MIP-CreER$^{TAM}$ mice were fed on HFD for one week and their pancreata was immunostained with Ki67 and insulin. β-cell proliferation was then analyzed. Data shown are the mean +/- SE (n=3-4, **p < 0.005).

**Supp. Figure 3. Nrf2 deletion decreases insulin content *ex vivo*.**
Isolated islets from NRF2$^{lox/lox}$ mice were transduced with LacZ or Cre expressing adenoviruses and incubated in 20 mM glucose. After 72 h islets were (A,B) immunolabeled with insulin and mean intensity was calculated, or (C) measured for insulin content, or (D) RNA was extracted and expression of β-cell identity genes was measured. Data shown are the mean +/- SEM (n=3-6, *p < 0.05; **p < 0.005, ****p < 0.0001).

**Supp. Figure 4. No differences of body weight or fasting blood glucose in βNrf2KO mice after 1 month on a HFD.** (A) Pancreata from βNrf2KO mice fed on HFD for 29 days and immunolabeled with antibodies against Nrf2 and insulin. (B) Fasting blood glucose and (C) body weight were measured in βNrf2KO mice fed on a HFD for

29 days. Data shown are the mean +/- SE (n=7-8, *p < 0.05; **p < 0.005; ****p < 0.0001).

**Supp. Figure 5. No changes in expression of maturation genes in Keap1$^{lox/lox}$ mouse β- cells.** Dispersed Keap1$^{lox/lox}$ islets were treated with Cre or LacZ expressing adenoviruses. Following 24 h, RNA was isolated and the expression of various β-cell identity genes was measured. Data shown are the mean +/- SE (n=3).

**Supp. Figure 6. β-cell-specific deletion of Keap1 *in vivo* does not result in changes of glucose homeostasis.** (A,B) Intraperitoneal glucose tolerance test (ipGTT) was performed in βKeap1KO mice after overnight fasting and the area under the curve (AUC) was calculated. (C,D) An insulin tolerance test (ITT) was performed in *ad lib* fed mice and the area under the curve (AUC) was calculated. (E) Fasting and (F) non-fasting blood-glucose were measured. (G) Body weight and (H) plasma insulin were measured. Data shown are the mean +/- SE (n=6-7).

**Supp. Figure 7. CDDO-Me increases Nrf2 levels and stimulate proliferation in INS-1 832/13 cells** (A) INS1 cells were incubated with the indicated concentrations of CDDO-Me for 72 h, followed by the addition of Trypan blue. Viable cells (unstained cells) were counted under a light microscope. (B) INS1 cells were incubated with 20 nM CDDO-Me for up to 50 min, followed by immunolabeling using an Nrf2-p antibody. Data shown are the mean +/- SE (n=3, *p < 0.05).

**Supplemental Figure 1**

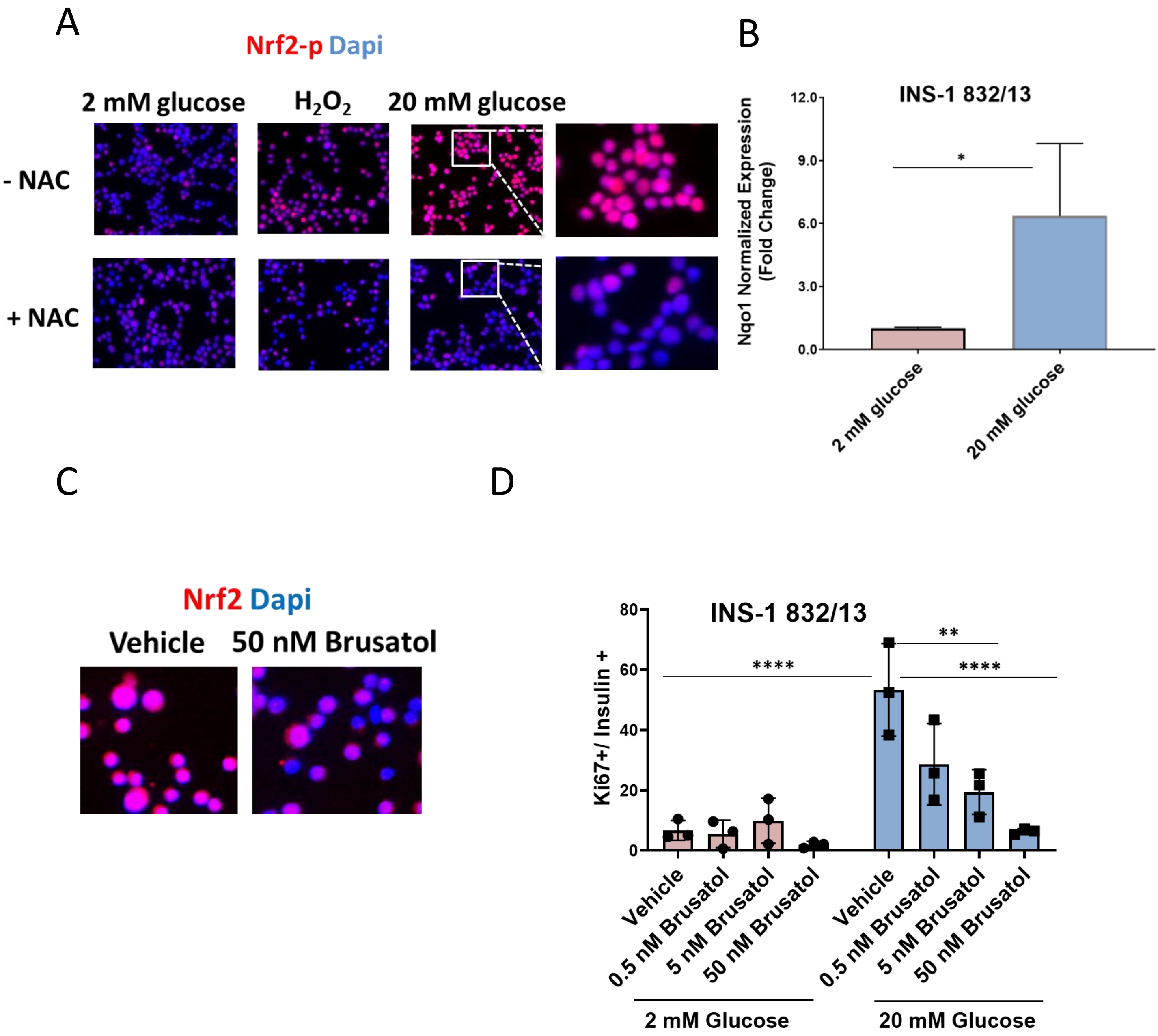

### Supplemental Figure 2

A

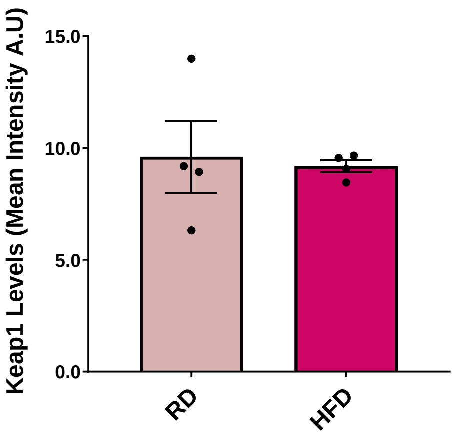


B

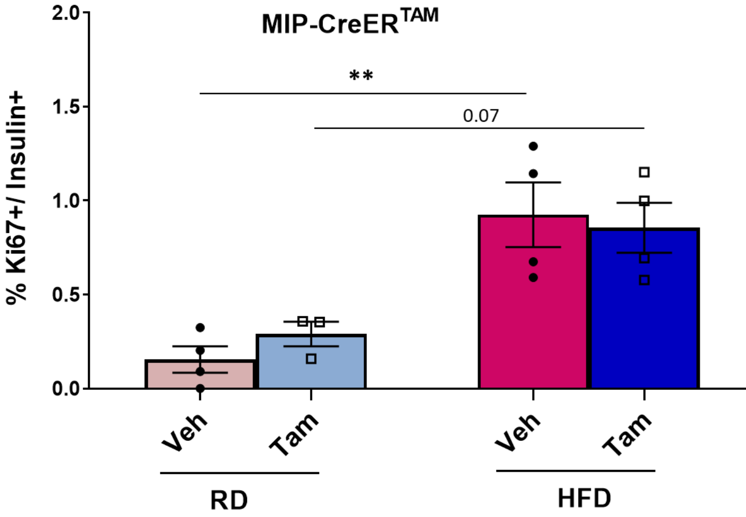

### Supplemental Figure 3

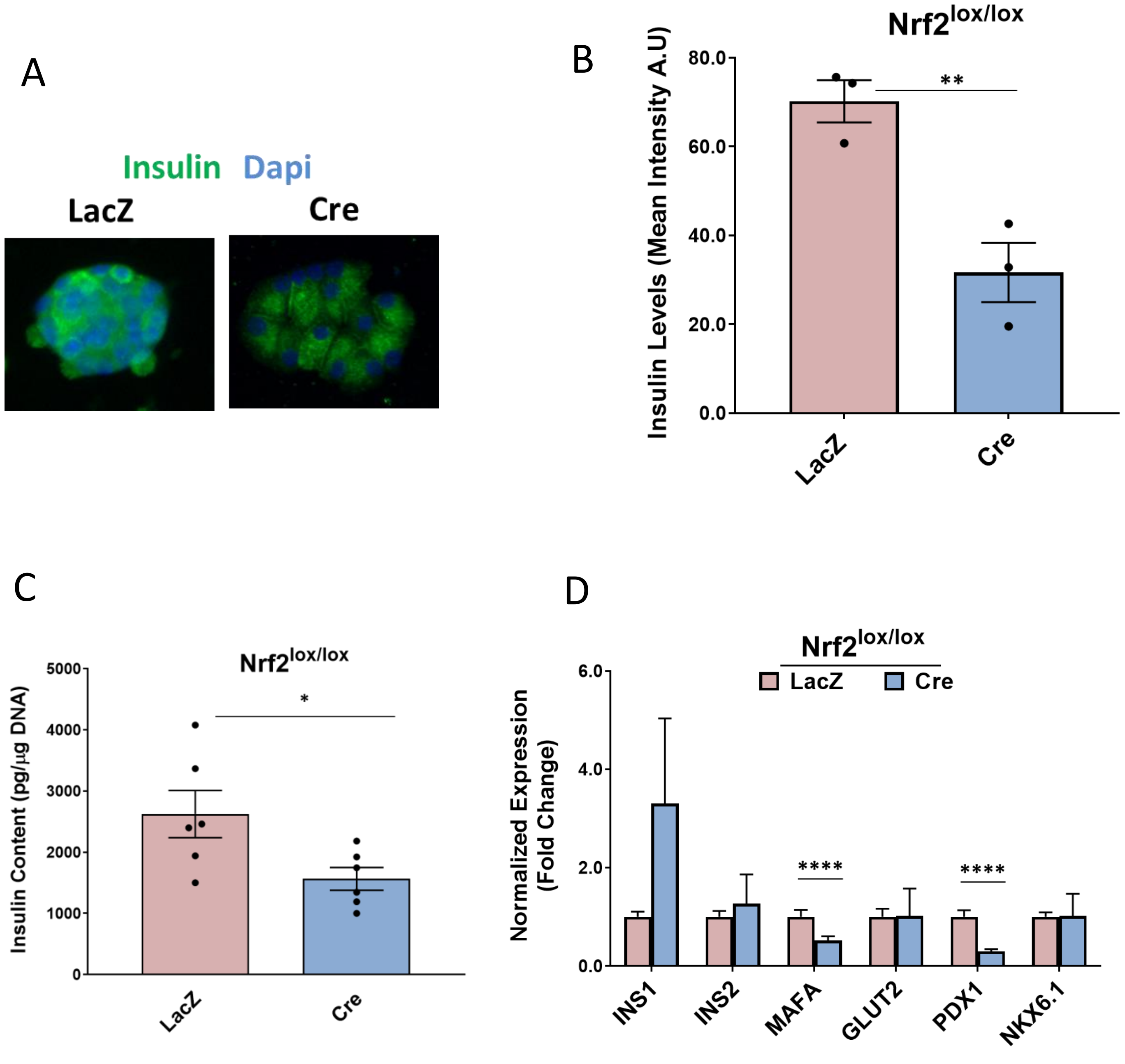

### Supplemental Figure 4

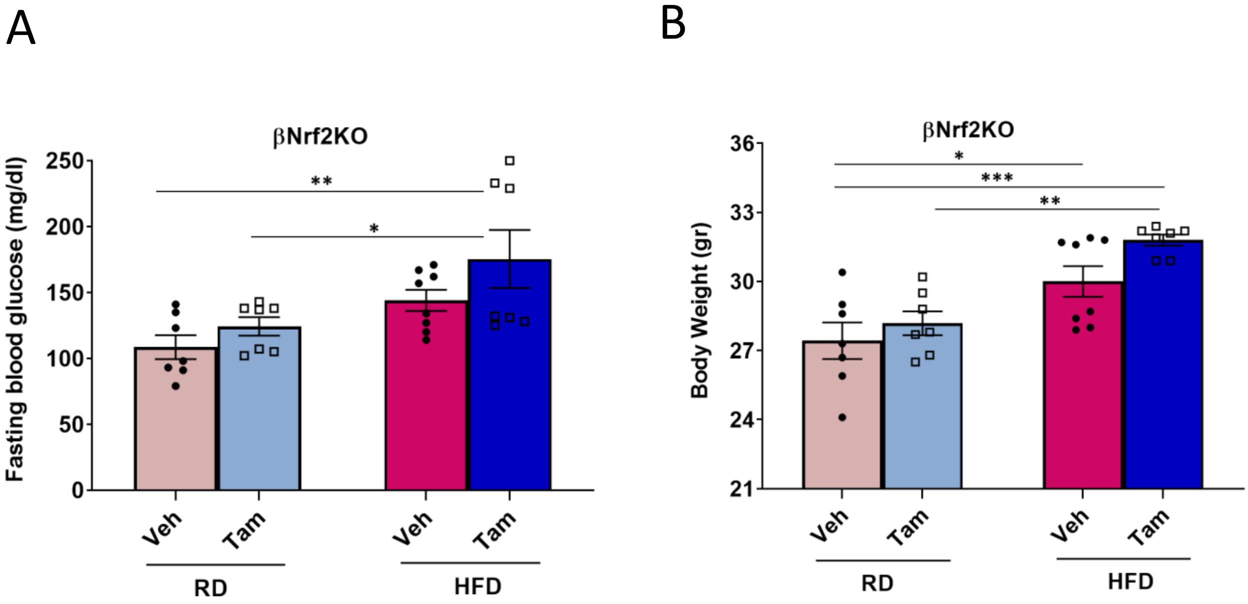

### Supplemental Figure 5

A

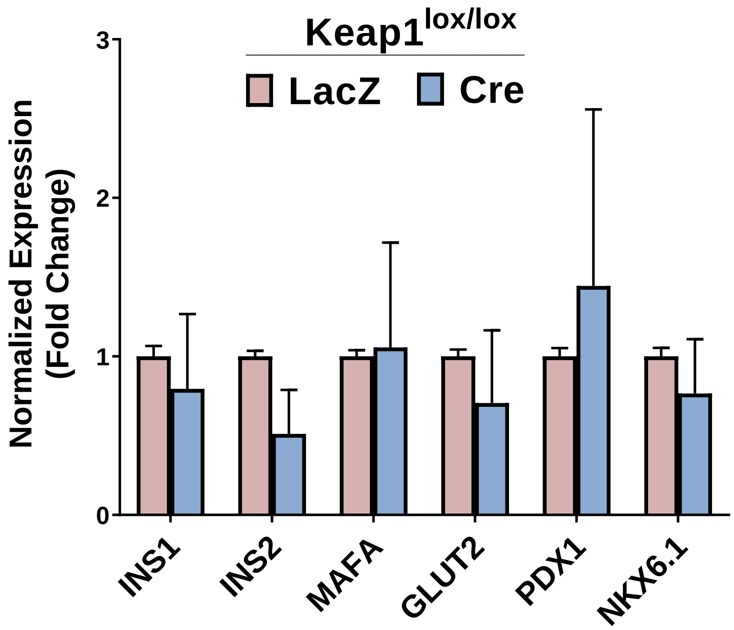

### Supplemental Figure 6

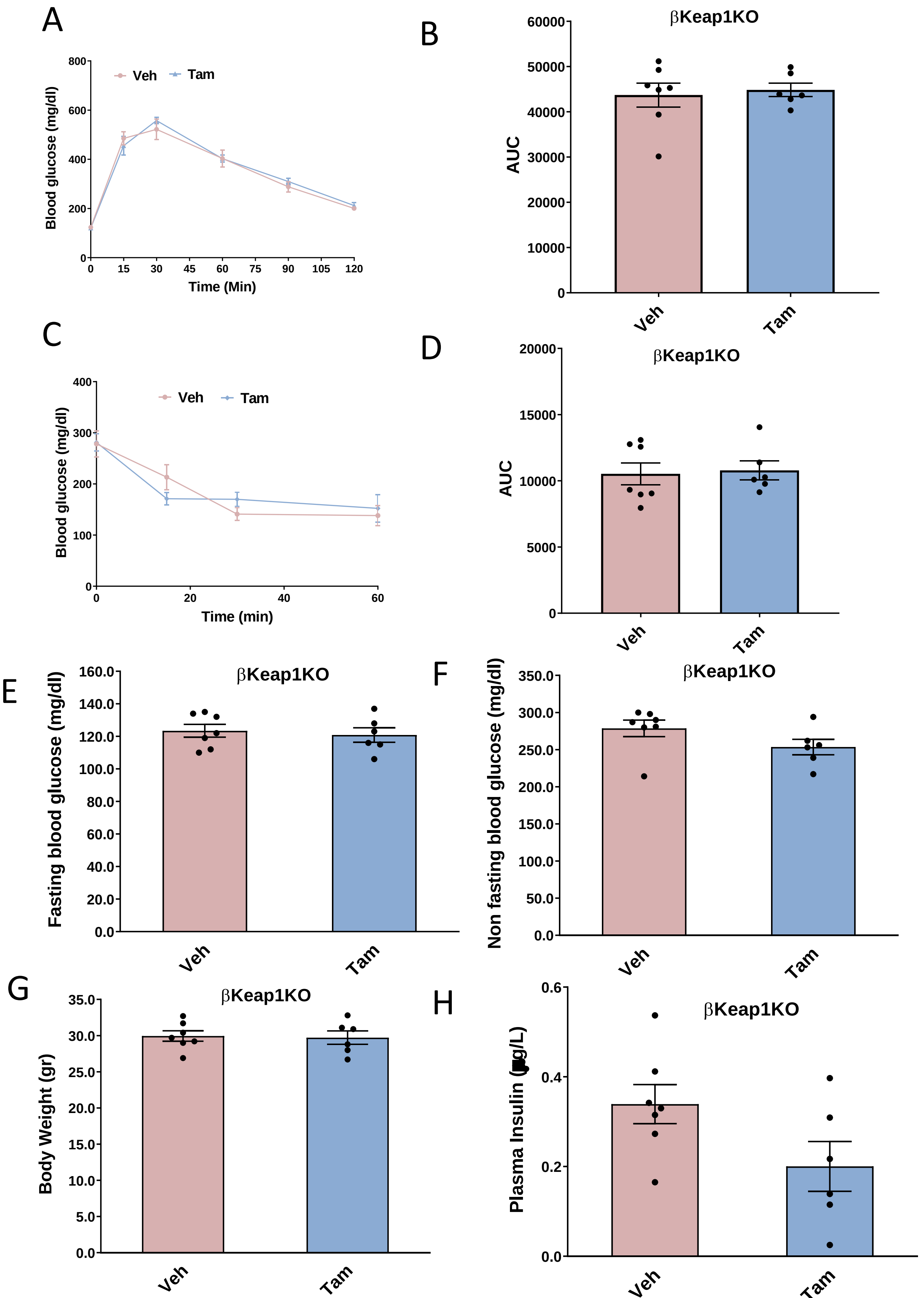

### Supplemental Figure 7

A

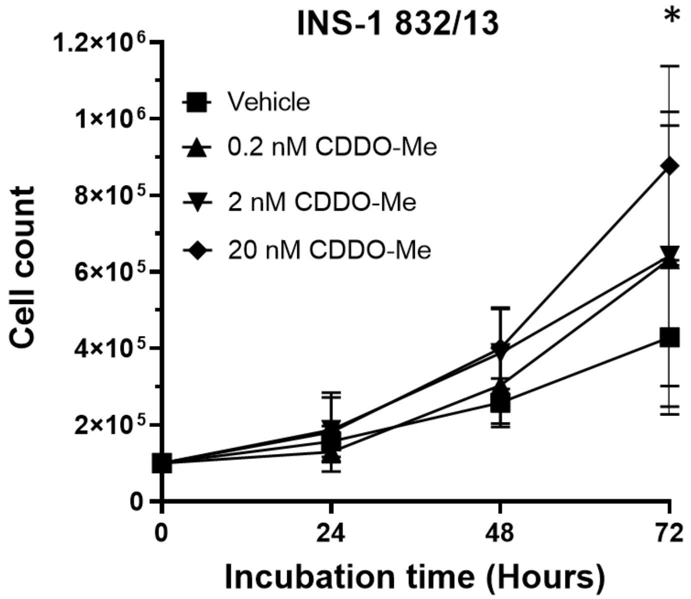


B

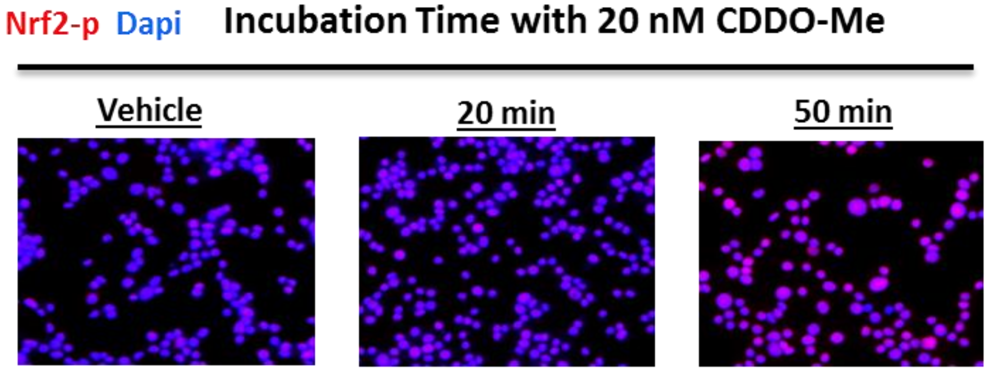
